# Natural variation in a single amino acid underlies cellular responses to topoisomerase II poisons

**DOI:** 10.1101/125567

**Authors:** Stefan Zdraljevic, Christine Strand, Hannah S. Seidel, Daniel E. Cook, John G. Doench, Erik C. Andersen

**Author notes:** Corresponding author Erik C. Andersen, Assistant Professor of Molecular Biosciences, Northwestern University, Evanston, IL 60208, USA.

## Abstract

Many medications, including chemotherapeutics, are differentially effective from one patient to the next. Understanding the causes of these population-wide differences is a critical step towards the development of personalized treatments and improvements to existing medications. Here, we investigate natural differences in sensitivity to anti-neoplastic drugs that target topoisomerase II, using the model organism *Caenorhabditis elegans*. We show that wild isolates of *C. elegans* vary in their sensitivity to these drugs, and we use an unbiased statistical and molecular genetics approach to demonstrate that this variation is explained by a methionine-to-glutamine substitution in topoisomerase II (TOP-2). The presence of a non-polar methionine at this residue increases hydrophobic interactions between TOP-2 and the poison etoposide, as compared to a polar glutamine. We hypothesize that this stabilizing interaction results in increased genomic instability in strains that contain a methionine residue. The residue affected by this substitution is conserved from yeast to humans and is one of the few differences between the two human topoisomerase II isoforms (methionine in hTOPIIα and glutamine in hTOPIIβ). We go on to show that this substitution influences binding and cytotoxicity of etoposide and two additional topoisomerase II poisons in human cell lines. These results explain why hTOPIIα and hTOPIIβ are differentially affected by various poisons and demonstrate the utility of *C. elegans* in understanding the genetics of drug responses.

## Introduction

Antineoplastic regimens used to treat cancer are often associated with poor prognoses and severe side effects. Ideally, antineoplastic regimens can be tailored to an individual patient based on various genetic markers known to be associated with drug response to maximize therapeutic effectiveness and minimize unwanted side effects. Advances in sequencing technologies over the course of the past decade promised the discovery of many genetic variants that contribute to human health. Though large-scale sequencing projects have lead to the identification of many genetic variants associated with disease risk (Visscher et al. 2012), relatively few variants have been identified that contribute to clinically relevant traits such as response to antineoplastic compounds. In fact, only 71 of over 500 FDA-approved antineoplastic compounds use genetic information to affect treatment efficacy (www.fda.gov). Unfortunately, the predictive power of these identified genetic variants can be inconsistent due to biases in the sampled population (Boddy 2013) and other key limitations of clinical genome-wide association (GWA) studies that attempt to link genetic variants with treatment outcomes. The major factor limiting the efficacy of these studies is sample size because it is difficult to identify large numbers of individuals exposed to the same antineoplastic regimens. This limitation is compounded when considering environmental (Liu et al. 2013; Hunter 2005) and tumor heterogeneity (Koboldt et al. 2013). As a result, most variants discovered to be associated with outcomes in clinical GWA studies offer low predictive power for patient responses to treatment (Park et al. 2012). These limitations and others emphasize the need for novel approaches to identify variants that predict patient outcomes to antineoplastic compounds.

Studies of model organisms have greatly facilitated our understanding of basic cellular processes. In recent years, *Saccharomyces cerevisiae* and *Drosophila melanogaster* have been used to understand the physiological effects of small molecules and repurposed as screening platforms to identify new antineoplastic compounds (Willoughby et al. 2013; Perlstein et al. 2007; King et al. 2014). The ability to generate extremely large numbers of recombinant yeast facilitates the identification of genomic regions that are predictive of drug response (Ehrenreich et al. 2010; Bloom et al. 2013). Furthermore, the specific genes and variants within regions can be identified and functionally validated in yeast (Demogines et al. 2008; Liti & Louis 2012; Stern 2014). By contrast, *D. melanogaster* studies offer the ability to study the physiological responses to drugs in the context of multiple tissue types, but functional validation of specific genes and variants associated with drug responses has been more limited (King et al. 2014). The roundworm *Caenorhabditis elegans* has the advantages of both S. *cerevisiae* and *D. melanogaster* because large cross populations can be generated to study the physiological responses to drugs in a metazoan. These attributes have made *C. elegans* an important model for connecting differential drug responses with genetic variants present in the species (Ghosh et al. 2012; Andersen et al. 2015).

Here, we take advantage of natural genetic variation present in *C. elegans* to identify the genetic basis underlying susceptibility to a panel of clinically relevant antineoplastic compounds that poison the activity of topoisomerase II enzymes. The inhibition of these enzymes by topoisomerase II poisons results in the accumulation of double-stranded breaks and genome instability (Pommier et al. 2010; Pommier et al. 1984; Gómez-Herreros et al. 2013). Topoisomerase II enzymes are targeted by antineoplastic regimens because proliferative cell populations require their enzymatic activity to relieve topological stress ahead of the replication fork (Nitiss 2009). Using two unbiased genetic mapping approaches, we show that divergent physiological responses to the topoisomerase II poison etoposide are determined by natural genetic variation in a *C. elegans* topoisomerase II enzyme. Furthermore, we show using CRISPR/Cas9-mediated genome editing that variation in a specific amino acid (Q797M) underlies the cytotoxic effects of etoposide. This residue is conserved in humans and is one of the few differences between the putative drug-binding pockets of the two topoisomerase II isoforms (M762 in hTOPIIα and Q778 in hTOPIIβ). Previous structural studies on hTOPIIβ implicated this glutamine residue in etoposide binding because of its proximity to the drug-binding pocket (Wu et al. 2011; Wu et al. 2013). However, a study on hTOPIIα suggested that the corresponding methionine residue has no functional role in drug binding (Wendorff et al. 2012). We present a mechanistic model to explain how variation at this residue underlies differential responses to etoposide and other topoisomerase II poisons. Finally, we use genome-edited human cell lines to show that this residue in hTOPIIα contributes to differential toxicity of various topoisomerase II poisons. These results demonstrate the power of using *C. elegans* natural genetic variation to identify mechanisms of drug susceptibility in human cells that could inform human health decisions based on genetic information.

## Results

### A single major-effect locus explains variation in response to etoposide

We investigated etoposide sensitivity in *C. elegans* using a high-throughput fitness assay. In brief, animals were grown in liquid culture in presence of etoposide, and body lengths of progeny and offspring production were measured using a COPAS BIOSORT (Supplemental figure 1). In this assay, shorter body lengths are indicative of developmental delay. To identify an appropriate dose of etoposide for this assay, we performed dose-response experiments on four genetically diverged isolates of *C. elegans:* N2 (Bristol), CB4856 (Hawaii), JU258, and DL238. We chose 250 μM etoposide for further experiments because it was the lowest concentration at which we observed an etoposide-specific effect in all four strains tested, trait differences between the laboratory Bristol strain (N2) and a wild strain from Hawaii (CB4856) strains were maximized, and the median animal length was highly heritable (Supplemental figure 2).

When grown in etoposide, progeny of the Hawaii strain are on average 75 μm shorter than progeny of the Bristol strain. To map the genetic variants underlying this difference, we performed our high-throughput fitness assay on a panel of 265 recombinant inbred advanced intercross lines (RIAILs), generated between a Bristol derivative (QX1430) and Hawaii (Andersen et al. 2015). We measured median animal length for each RIAIL strain grown in etoposide, and we corrected for assay-to-assay variability and effects of the drug carrier (DMSO) using a linear model. We used the resulting regressed median animal length trait (referred to as animal length) for quantitative trait locus (QTL) mapping. This mapping identified a major-effect QTL for etoposide resistance on chromosome II at 11.83 Mb (Figure 1A). This QTL explained 27% of the phenotypic variance among the recombinant lines. The QTL confidence interval spans from 11.67 to 11.91 Mb on chromosome II and contains 90 genes, 68 of which contain variation between the parental strains.

**Figure 1.**
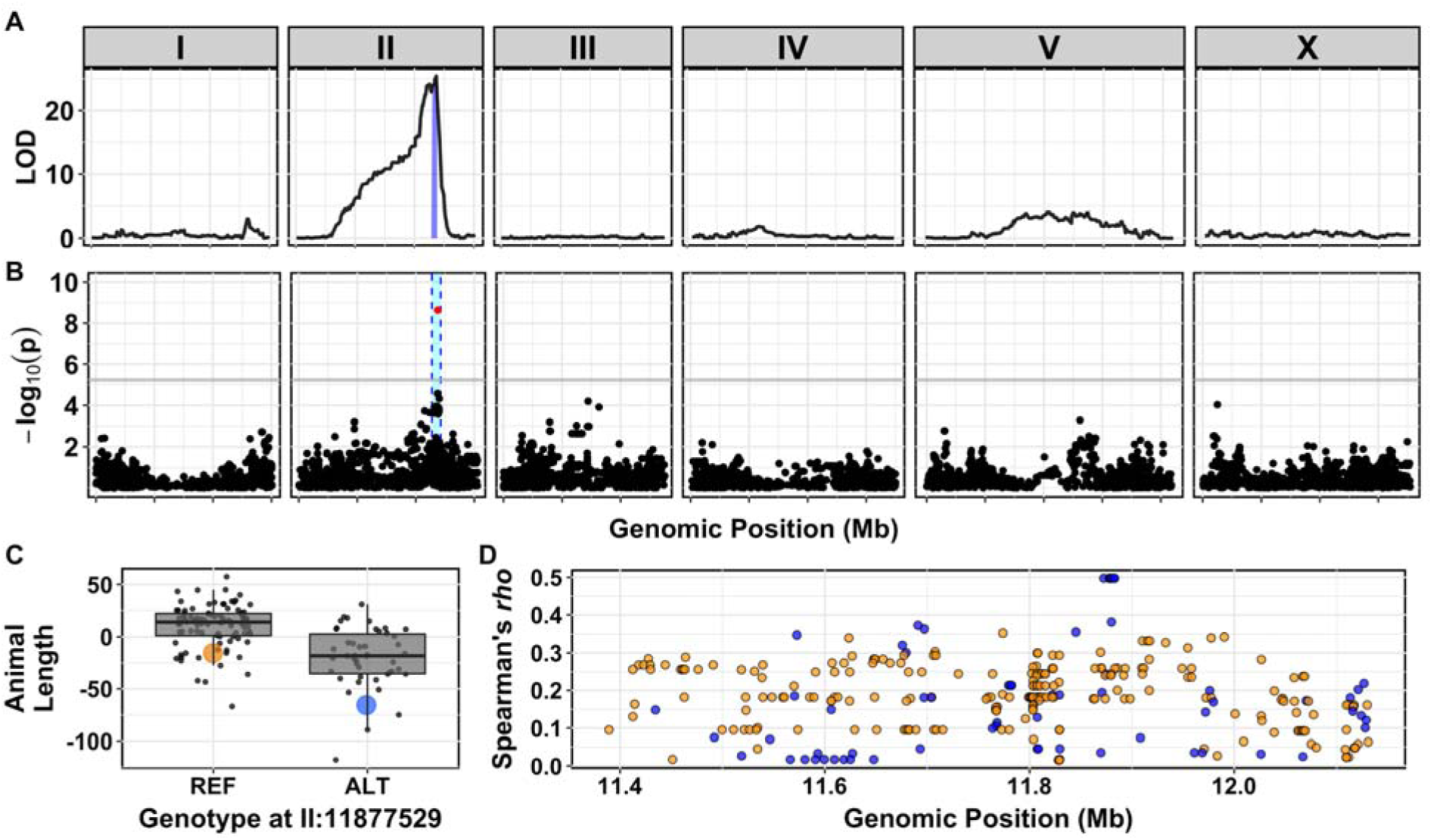
A major-effect locus controls differences in etoposide response. (A) Linkage mapping plot for regressed median animal length in the presence of etoposide is shown. The significance value (logarithm of odds, LOD, ratio) is plotted on the y-axis, and genomic position (Mb) separated by chromosome is plotted on the x-axis. Each tick on the x-axis corresponds to 5 Mb. The associated 1.5 LOD-drop confidence intervals are represented by blue bars. (B) A manhattan plot for regressed median animal size in the presence of etoposide is shown. Each dot represents an SNV that is present in at least 5% of the phenotyped population. The −*log*_10_(*p*) for each SNV is plotted on the y-axis, and the genomic position (Mb) is plotted on the x-axis. Each tick on the x-axis corresponds to 5 Mb. SNVs are colored red if they pass the genome-wide Bonferroni-corrected significance threshold, which is denoted by the gray horizontal line. Genomic regions of interest are represented by a cyan rectangle. (C) Tukey box plots of phenotypes used for association mapping in (B) are shown. Each dot corresponds to the phenotype of an individual strain, which is plotted on the y-axis. Strains are grouped by their genotype at the peak QTL position, where REF corresponds to the allele from the reference Bristol strain. The Bristol and Hawaii strains are colored orange and blue, respectively. (D) Fine mapping of the chromosome II region of interest, showing the calculated Spearman’s *rho* statistic for every variant present in the QTL confidence interval from (A), is shown. The genomic position of each SNV is on the x-axis in Mb. Cyan dots correspond to variants that are present in the Hawaiian strain, and gray dots correspond to variants that are present in the wild population.

We next sought to validate this QTL using homozygous reciprocal near-isogenic lines (NILs), which contain either the QTL confidence interval from the Bristol strain introgressed into the Hawaii strain or the interval from the Hawaii strain introgressed into the Bristol strain. NILs with the genomic interval derived from the Bristol strain have increased resistance to etoposide compared to the Hawaii strain (Supplemental figure 3). Similarly, NILs with the genomic interval derived from the Hawaii strain exhibited decreased resistance to etoposide. These results confirmed that genetic variation located on the right arm of chromosome II contributes to differential etoposide susceptibility.

### The same locus on chromosome II explains variation in response to etoposide in a panel of wild *C. elegans* isolates

In the initial dose response experiments, we found that JU258 and DL238 had different responses to etoposide than the Bristol and Hawaii strains, suggesting that additional genetic variation present in the wild *C. elegans* population could also contribute to etoposide response. To identify this additional variation, we performed a genome-wide association (GWA) mapping of etoposide resistance in 138 wild *C. elegans* isolates. This analysis led to the identification of a QTL on the right arm of chromosome II with a peak position at 11.88 Mb (Figure 1B). This QTL has a genomic region of interest that spans from 11.70 to 12.15 Mb for which we found no evidence of selection (Supplemental figure 4) or geographic clustering of the peak QTL allele (Supplemental figure 5). In addition, this QTL overlaps with the QTL identified through linkage mapping described above. Of the 138 wild isolates assayed, including the Hawaiian strain, 46 have the alternate (non-Bristol) genotype at the peak position on chromosome II (Figure 1C). Similar to our observations using the recombinant lines, the 46 strains that contain the alternate genotype are more sensitive to etoposide than strains containing the Bristol genotype at the QTL peak marker. We hypothesized that variation shared between the Hawaiian strain and the other 45 alternate-genotype strains contributes to etoposide sensitivity because we detected overlapping QTL, with the same direction of effect, between GWA and linkage mapping experiments. This hypothesis suggested that we could condition a fine-mapping approach on variants found in the Hawaiian strain and shared across these 45 strains.

To fine-map the QTL, we focused on variants shared among wild isolates. Using data from the *C. elegans* whole-genome variation dataset (Cook, Zdraljevic, Tanny, et al. 2016) we calculated Spearman’s *rho* correlations between animal length and each single-nucleotide variant (SNV) in the QTL confidence interval (Figure 1D). SNVs in only three genes, *npp-3, top-*2, and *ZK930.5*, were highly correlated with the etoposide response (*rho* > 0.45). Of these genes, the *top-2* gene encodes a topoisomerase II enzyme that is homologous to the two human isoforms of topoisomerase II. We prioritized *top-2* because topoisomerase II enzymes are the cellular targets for etoposide (Pommier et al. 2010).

### Genetic variation in *top-2* contributes to differential etoposide sensitivity

To determine if genetic variation present in the *top-2* gene contributes to differential etoposide sensitivity, we performed a reciprocal hemizygosity test (Stern 2014). Prior to this test, we determined that resistance to etoposide is dominant by measuring the lengths of F1 heterozygotes from a cross between the Bristol and Hawaii strains in the presence of etoposide (Supplemental figure 6). Additionally, we tested *npp-3* and *top-2* deletion alleles from the Bristol genetic background and found that only loss of *top-2* contributes to etoposide sensitivity (Supplemental figure 7). To more definitively show a causal connection of *top-2* variation to etoposide sensitivity, we used a reciprocal hemizygosity test. First, we introgressed the *top-2*(*ok1930*) deletion allele into the Hawaiian genetic background. The Bristol/Hawaii(*Δtop-2*) heterozygote that contains the Bristol *top-2* allele is more resistant to etoposide treatment than the Hawaii/Bristol(*Δtop-2*) heterozygote, which suggests that the Bristol *top-2* allele underlies etoposide resistance (Figure 2A, Supplemental figure 8). The observed differences between the Hawaii/Bristol(*Δtop-2*) and Bristol/Hawaii(*Δtop-2*) heterozygotes confirmed that *top-2* variation underlies differential susceptibility to etoposide.

**Figure 2.**
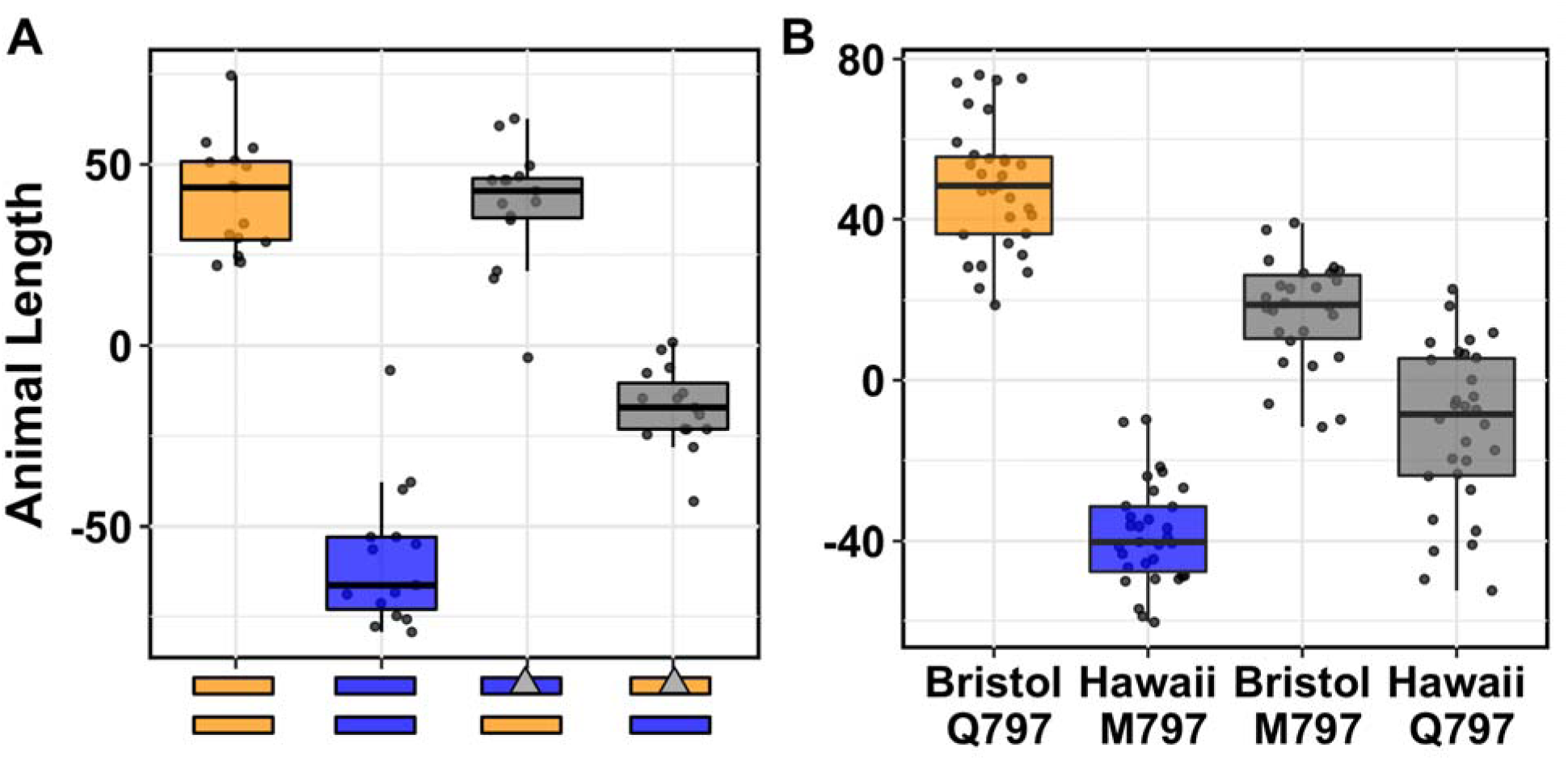
Q797M variant in TOP-2 underlies etoposide sensitivity in *C. elegans*. (A) Tukey box plots of the residual median animal length distribution of Bristol (orange) and Hawaii (blue) compared to two heterozygous *top-2* deletion strains (gray) in response to etoposide are shown. Orange (Bristol) and blue (Hawaii) rectangles below the plot correspond to the two chromosome II homolog genotypes. Gray triangles denote chromosomes with the *top-2* deletion allele. The Bristol strain and the Bristol/Hawaii(Δ*top-2*) heterozygous strain are not significantly different from each other (Tukey HSD, p-value 0.9990812), but all other comparisons are significant (Tukey HSD p-value < 0.0001). (B) Tukey box plots of residual median animal length after etoposide exposure are shown (Bristol, orange; Hawaii, blue; allele replacement strains, gray). Labels correspond to the genetic background and the corresponding residue at position 797 of TOP-2 (Q for glutamine, M for methionine). Every pair-wise strain comparison is significant (Tukey HSD, *p*-value < 2.0E-6).

### A glutamine-to-methionine variant in TOP-2 contributes to etoposide response

To identify genetic variants in *top-2* that contribute to etoposide resistance in the Bristol strain, we focused on genomic differences between the Bristol and Hawaii strains. Based on gene expression data between the Bristol and Hawaii strains (Rockman et al. 2010), *top-2* is expressed at similar levels. Therefore, we concluded that etoposide resistance in the Bristol strain is likely caused by coding variation. The *C. elegans top-2* gene contains 31 SNVs across the population-wide sample of 138 wild isolates. We narrowed our search to 16 variants present in the Hawaiian strain. Two of these variants are in the 3’ UTR, three are in introns, and six are synonymous variants that likely do not contribute to etoposide resistance. The remaining five variants encode for amino acid changes in the TOP-2 enzyme. Of these five variants, four were highly correlated with etoposide sensitivity in the wild isolate panel: Q797M, I1206L, Q1217A, and D1387N. Multiple-sequence alignment of TOP-2 peptides across yeast, *D. melanogaster*, mice, and humans revealed that I1206L, Q1217A, and D1387N are in the variable C-terminal domain (Supplemental file 1). By contrast, the Q797M variant is located in the conserved DNA binding and cleavage domain (Schmidt et al. 2012). Structural data suggest that the TOP-2 Q797 residue lies within the putative etoposide-binding pocket (Wu et al. 2011), and the corresponding residue is a methionine (M762) in the hTOPOIIα and a glutamine (Q778) in hTOPOIIβ (Wendorff et al. 2012). Additionally, the two human isoforms differ in one other residue within the putative etoposide-binding pocket (S800(α)/A816(β)). Therefore, the *C. elegans* glutamine-to-methionine TOP-2 variant mirrors one of two differences within the etoposide-binding pocket of the two human topoisomerase II enzyme isoforms. Crucially, hTOPOIIα forms a more stable DNA-TOPOII cleavage complex with etoposide than hTOPOIIβ (Bandele & Osheroff 2008). We hypothesized that etoposide sensitivity in both *C. elegans* and the human isoforms is affected by this residue.

To test the effects of the Q797M variant on *C. elegans* response to etoposide, we used CRISPR/Cas9-mediated genome editing to change this residue. We replaced the glutamine residue in the Bristol strain with a methionine and the methionine residue in the Hawaii strain with a glutamine. We exposed the allele-replacement strains to etoposide and found that the methionine-containing Bristol animals were more sensitive than glutamine-containing Bristol animals (Figure 2B). Conversely, the glutamine-containing Hawaii animals were more resistant to etoposide than the methionine-containing Hawaii animals (Figure 2B). These results confirm that this variant contributes to differential etoposide sensitivity between the Bristol and Hawaii strains.

### Methionine mediates stronger hydrophobic interactions with etoposide than glutamine

We hypothesized that the non-polar functional group attached to the glycosidic bond of etoposide contributes to increased stability of the drug-enzyme complex by forming a more stable interaction with the methionine residue than with the glutamine residue. To test this hypothesis, we simulated etoposide docking into the putative drug-binding pocket of the TOP-2 homology model generated by threading the *C. elegans* peptide sequence into the hTOPOIIβ structure (RMSD = 1.564Å, PDB:3QX3; (Wu et al. 2011)). Upon etoposide binding, the free energy (ΔG) of the drug-binding pocket was -10.09 Kcal/mol for TOP-2 Q797 (Figure 3A) and - 12.67 Kcal/mol for TOP-2 M797 (Figure 3B). This result suggests that etoposide interacts more favorably with TOP-2 M797 than with TOP-2 Q797, consistent with our results in live animals. A more favorable drug-enzyme interaction, as indicated by a more negative ΔG, likely causes increased stability of the TOP2 cleavage complexes, which has been shown to result in a greater number of double-stranded breaks throughout the genome (Deweese & Osheroff 2009). Therefore, we expect *C. elegans* strains that contain a methionine at this residue to accumulate more genomic damage when exposed to etoposide. The resulting physiological effect of increased genomic damage likely delays development and causes the progeny of exposed individuals to be shorter.

**Figure 3.**
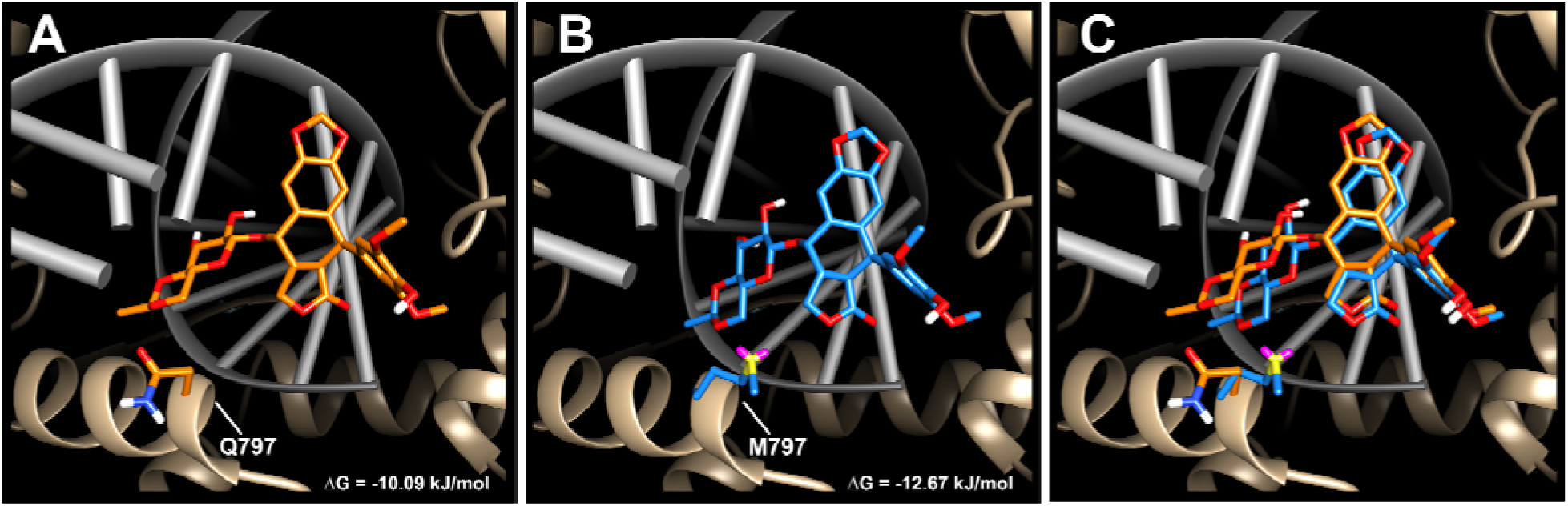
Etoposide docked into the *C. elegans* TOP-2 homology model. Etoposide docked into the (A) glutamine-containing Bristol TOP-2 enzyme (ΔG = -10.09 kJ/mol), (B) methionine-containing Hawaii TOP-2 enzyme (ΔG = -12.67 kJ/mol), and (C) the overlay of both structures. Glutamine is colored in orange, and methionine is colored in blue. Etoposide docked into the glutamine-containing TOP-2 enzyme is orange, and etoposide docked into the methionine-containing TOP-2 enzyme is blue. DNA is colored in gray, and the ribbon representation of the TOP-2 protein is shown in tan.

### TOP-2 variation causes allele-specific interactions with an expanded set of topoisomerase II poisons

Because the molecular docking simulations explain the observed physiological effects of etoposide exposure, we hypothesized that the 797 residue of TOP-2 would mediate differential interactions with additional topoisomerase II poisons based on their chemical structures. Like etoposide, teniposide, dactinomycin, and amsacrine each contain core cyclic rings that are thought to interfere with the re-ligation step of the topoisomerase II catalytic cycle through DNA interactions (Pommier et al. 2010). However, the functional groups attached to the core cyclic rings of each poison vary in their polarity and size, which could affect interactions with topoisomerase II enzymes. For example, the only difference between teniposide and etoposide is the presence of a thienyl or methyl group attached to the D-glucose derivative, respectively, but they share a similarly sized and hydrophobic functional group. We predicted that these two drugs would have comparable interactions with the TOP-2 alleles and elicit a similar physiological response. By contrast, the polar functional groups of dactinomycin likely have stronger interactions with the glutamine variant and induce increased cytotoxicity in animals that contain this allele. We quantified the physiological responses of the TOP-2 allele-replacement strains exposed to these two drugs and found that each response matched our predictions (Figure 4). Specifically, strains harboring the TOP-2 methionine allele were more sensitive to teniposide than those strains that contain the glutamine allele. Conversely, strains with the TOP-2 glutamine allele were more sensitive to dactinomycin than those strains with the methionine allele. Unlike etoposide, teniposide, or dactinomycin, the core cyclic rings of amsacrine do not have an equivalent functional group to interact with the TOP-2 797 residue, suggesting that variation at TOP-2 residue 797 will have no impact on amsacrine sensitivity. Although the Bristol and Hawaiian strains differed, we found that the allele status of TOP-2 had no quantifiable effect on amsacrine response (Figure 4) and different genomic loci control response to this drug (Supplemental figure 9). These results support the hypothesis that the polarity of the putative drug-binding pocket determines the cytotoxic effects of multiple, but not all, topoisomerase II poisons. To further explore this hypothesis, we tested a drug (XK469) that has preferential hTOPOIIβ specificity (Gao et al. 1999). Surprisingly, we found that the strains that contain the methionine allele (like hTOPOIIα) were more sensitive to XK469 (Supplemental figure 10). This result indicates that an additional mechanism might contribute to XK469 specificity in human cells and underscores the importance of functional validation of specific residues that are thought to be involved in targeted drug binding.

**Figure 4.**
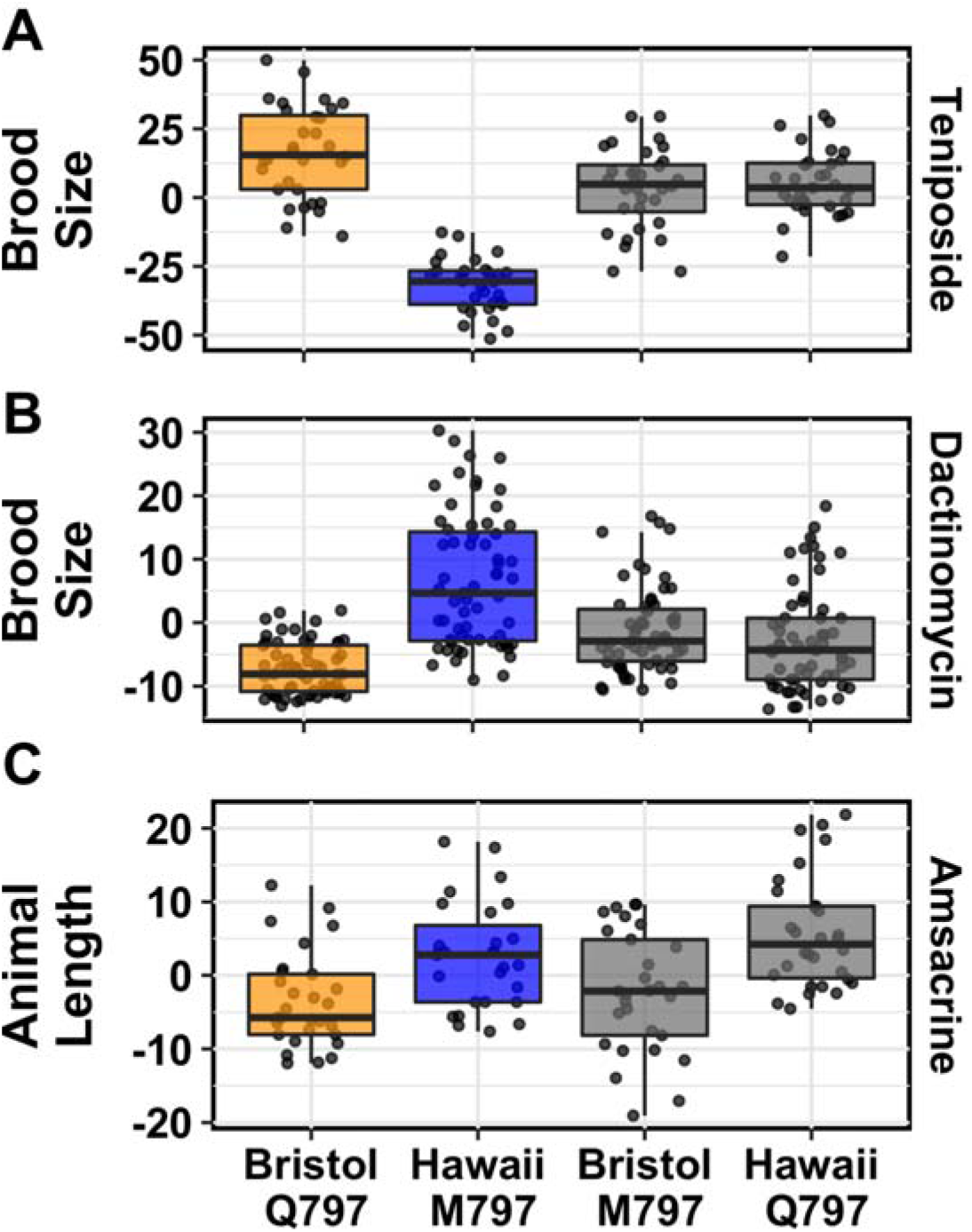
Variation in the TOP-2 797 allele underlies response to multiple topoisomerase II poisons. Tukey box plots of (A) regressed brood size in response to teniposide show that the strains containing the glutamine allele of TOP-2 are significantly more resistant to teniposide than strains with the methionine allele (Tukey HSD, *p*-value < 1E-16). (B) Conversely, Tukey box plots of regressed brood size in response to dactinomycin show that strains with the methionine allele are significantly more resistant to dactinomycin than those with with the glutamine allele (Tukey HSD, *p*-value = 2.0E-4). (C) By contrast, Tukey box plots of regressed median animal length in response to amsacrine show that the TOP-2 797 residue had no effect on amsacrine response (Tukey HSD, *p*-value > 0.414). Orange corresponds to the Bristol genetic background and blue to the Hawaii background. Labels correspond to the genetic background and the corresponding residue at position 797 of TOP-2 (Q for glutamine, M for methionine).

### Variation in the equivalent site in topoisomerase II alpha causes differential susceptibility to diverse poisons in human cells

To determine if differences in the hydrophobicities of the two human topoisomerase II putative drug-binding pockets underlie etoposide sensitivity, we used CRISPR/Cas9 genome editing and a pooled-sequencing approach to create human embryonic kidney 293 cells (293T) that encode hTOPOIIα enzymes with a hTOPOIIβ-like drug-binding pocket. Cells were incubated with genome-editing machinery for six hours, allowed to recover for five days, and then split into two populations for etoposide exposure or no etoposide exposure. Etoposide treatment provided a selective pressure that upon further passaging led to a greater than 160-fold enrichment of cells that contain the glutamine-edited hTOPOIIα allele as compared to populations of cells exposed to no drug (Figure 5). These results show that cells with the glutamine-edited hTOPOIIα allele are more resistant to etoposide treatment than cells with the non-edited methionine hTOPOIIα allele. Notably, the rarity of genome editing events makes it unlikely that every copy of the hTOPOIIα gene in this diploid/polyploid cell line is edited. Because we see etoposide resistance in these incompletely edited cells, hTOPOIIα dimeric complexes likely contain one edited and one wild-type copy of hTOPOIIα and do not bind etoposide as well as causing less cytotoxicity. These data confirm both our dominance test (Supplemental figure 6) and the two-drug model of etoposide binding (Bromberg et al. 2003) in which both enzymes of the homodimer must be bound by poison to be completely inhibited. Additionally, we performed the reciprocal experiment to edit the glutamine-encoding hTOPOIIβ gene to a version that encodes methionine. If the methionine hTOPOIIβ allele is more sensitive to etoposide than the glutamine hTOPOIIβ, we would expect to observe a depletion of methionine-edited cells upon etoposide treatment. However, because glutamine-to-methionine editing occurred in less than 1% of the cells, it was difficult to detect further reductions in methionine allele frequencies (Supplemental table 1). Overall, we demonstrate that this residue underlies variation in etoposide response in both *C. elegans* and human cell lines.

**Figure 5.**
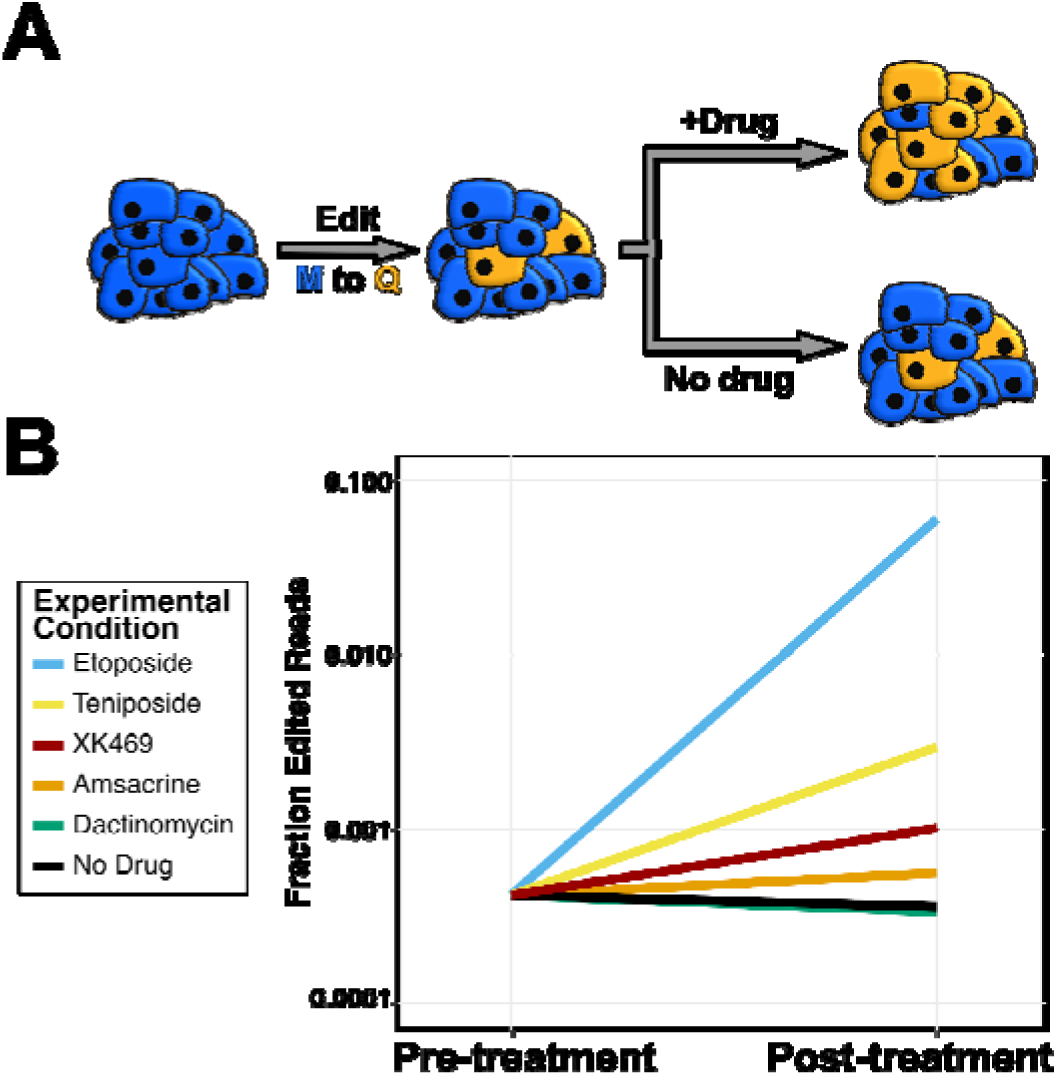
Human cells that contain hTOPOIIα M762Q are more resistant to etoposide, teniposide, and XK469. (A) Cartoon depiction of human cell-line experiment. A population of cells is incubated with CRISPR reagents to introduce the M762Q mutation in hTOPOIIα. A small fraction of cells are edited at this position and depicted in orange. Prior to splitting this population of cells into separate growth cultures with drug or control conditions, a subset of cells are prepared for sequencing to assess the fraction of edited cells. Split populations are then allowed to grow for two weeks and prepared for sequencing. In this example, blue cells contain the wt hTOPOIIα and the orange cells contain the edited M762Q hTOPOIIα, both strains contain the wt hTOPOIIβ. (B) Log-transformed fraction of sequencing reads that contain the CRISPR-edited allele of hTOPOIIα is plotted on the y axis for pre-treatment and post-treatment cell populations. The fraction of cells that contain the hTOPOIIα glutamine allele increase by 168.2-, 8.2-, and 2.8-fold upon treatment with etoposide, teniposide, or XK469, respectively, when compared to the no-drug control.

Our *C. elegans* results using the genome-edited *top-2* strains show that variation at this residue underlies differences in some but not all topoisomerase II poisons. We exposed the edited human cell lines to these different poisons to test this hypothesis. We found that cells containing the glutamine-edited hTOPOIIα allele are less affected by both teniposide and XK469, as indicated by the respective 8.2- and 2.8-fold increase in edited cell frequency upon drug exposure as compared to cell populations with no drug added. These results mirror our observations that *C. elegans* strains with the methionine TOP-2 allele are more sensitive to these drugs. We observed moderate-to-no change in edited allele frequencies between cell populations exposed to amsacrine (∼1.6-fold increase) or dactinomycin (∼0.93-fold decrease) and those cells exposed to no-drug control conditions. These results indicate that the hTOPOIIα M762 residue does not interact with amsacrine. Our expectation was that dactinomycin would be more cytotoxic to cells that contain the glutamine hTOPOIIα allele, and therefore result in a depletion of edited cells. As mentioned above, we do not have the power to detect this depletion in response because of low CRISPR-editing efficiency. However, we did expect to see enrichment of hTOPOIIβ Q778M edited cells upon exposure to dactinomycin in the reciprocal experiment. Despite this expectation, we saw no change in hTOPOIIβ Q778M edited cell frequency between dactinomycin and the no drug control. Our inability to detect any enrichment of allele frequencies might result from dactinomycin having multiple cellular targets in addition to topoisomerase II (Koba & Konopa 2005). Taken together, our findings testing a variety of poisons on human cell lines recapitulated the results from *C. elegans*.

## Discussion

Few genetic markers have been identified that predict patient responses to chemotherapeutic regimens (Moen et al. 2012; Giacomini et al. 2007). The goal of this study was to introduce new methods for the rapid and cost-effective identification of genetic variants that explain differences in chemotherapeutic response. Our approach leveraged genetic and phenotypic variation present in the model organism *C. elegans* to identify a single amino acid variant (Q797M) in the topoisomerase II enzyme that underlies differences in etoposide response. Mechanistic insights into differential etoposide binding between the glutamine or methionine alleles gave us the power to predict the physiological responses to an expanded panel of topoisomerase II poisons. These results highlight how the combination of a highly sensitive phenotyping assay with classical and quantitative genetics approaches in *C. elegans* can rapidly identify the mechanistic underpinnings of phenotypic variability in response to a key class of antineoplastic compounds.

Our approach stands in stark contrast to previous underpowered human cell line (Huang et al. 2007) and clinical studies (Low et al. 2013) that failed to identify any statistically significant associations between etoposide-induced cytotoxicity and genetic variation in the human population. However, the residue we identified in *C. elegans* does not vary in the human population (Lek et al. 2016), suggesting that GWA studies would not have identified this variant as a marker for etoposide sensitivity. Nevertheless, this residue is one of the few differences between the putative drug-binding pockets of the two human topoisomerase II isoforms (M762 in hTOPIIα and Q778 in hTOPIIβ), which allowed us to investigate the molecular underpinnings of drug binding. We verified that this single amino acid change in the human topoisomerase II isoforms results in profound differences in topoisomerase II poison-induced cytotoxicity using 293T cells. Though previous hTOPOIIβ structural studies have implicated this glutamine-methionine difference as functionally important for etoposide binding (Wu et al. 2011; Wu et al.2013), studies involving hTOPOIIα have argued that this residue is not involved (Wendorff et al.2012). The results presented here unequivocally show that this residue contributes to differential topoisomerase II poison-induced cytotoxicity and have important implications for targeted drug design.

Although topoisomerase II poisons can bind and inhibit both hTOPOIIα and hTOPOIIβ, hTOPOIIα is the cellular target of poisons in most cancers because it is expressed in proliferating cells (Pommier et al. 2010). However, recent evidence suggests that side effects associated with these treatments are caused by inhibition of hTOPOIIβ in differentiated cells (Chen et al. 2013). For example, antineoplastic treatment regimens that contain the epipodophyllotoxins (*e.g*. etoposide or teniposide) are hypothesized to increase the risk of developing secondary malignancies caused by hTOPOIIβ-dependent 11q23 translocations (Felix et al. 2006; Cowell et al. 2012; Azarova et al. 2007; Ratain et al. 1987). Additionally, the most severe side effects associated with treatments that contain an alternative class of topoisomerase II poisons (anthracyclines, *e.g*. doxorubicin or daunorubicin) include dose-dependent cardiotoxicity and heart failure dependent on hTOPOIIβ (Zhang et al. 2012; Vejpongsa & Yeh 2014; Yeh & Bickford 2009). Therefore, optimal topoisomerase II poisons will maximize interactions with hTOPOIIα to inhibit proliferating cells and minimize hTOPOIIβ interactions to reduce side effects. With this goal in mind, others have identified etoposide analogues with different isoform specificities but have not determined the mechanism of specificity (Mariani et al. 2015). Our study functionally validates a key residue determining isoform specificity and is critical to the improvement of this widely administered drug class. The importance of such functional characterization is underscored by our observation that *C. elegans* strains and human cells with the methionine TOP-2 allele are more sensitive to XK469, despite this drug being shown to be a β-specific poison (Gao et al. 1999). Though XK469 has been shown to be a β-specific poison, no information regarding its drug binding pocket or the mechanism driving isoform specificity is currently known. Our results indicate that XK469 occupies a similar drug-binding pocket of TOP-2 as other topoisomerase II poisons and interacts with residue 797.

To date, no human genetic variants have been linked to topoisomerase II poison-induced cytotoxicity. Of the 291 and 279 respective SNVs in hTOPOIIα and hTOPOIIβ that encode for missense mutations (Supplemental table 2 and 3), some are near the highly conserved DNA-binding domains or drug-binding pockets (Supplemental figure 11), which could affect drug response. However, the extent to which these variants impact responses to topoisomerase II poisons is unknown, so functional validation is required. The approach of editing human cells and following allele frequencies via sequencing represents a scalable method to assess the functional role of these variants and avoids single-cell cloning. Importantly, differences in responses to topoisomerase II poisons might not be affected by variation in the topoisomerase II isoforms but instead mediated by variation in cellular import, metabolism, or export. Pharmacogenomic data available for many antineoplastic compounds (Yang et al. 2009; Sim et al. 2011), in combination with human variation data (Lek et al. 2016), can be used to prioritize and test variants in highly conserved regions of proteins known to be involved in these alternative processes. This biased approach focused on candidate variants is necessitated by the lack of power in clinical GWA studies and is not guaranteed to successfully connect variants to differences in drug response. For this reason, unbiased mapping approaches in model organisms combined with functional validation in genome-edited human cells will greatly expand our current understanding of how human genetic variation affects drug responses.

## Experimental Procedures

### Strains

Animals were cultured at 20°C with the bacterial strain OP50 on modified nematode growth medium (NGM), containing 1% agar and 0.7% agarose to prevent burrowing of the wild isolates. For each assay, strains were grown at least five generations with no strain entering starvation or encountering dauer-inducing conditions (Andersen et al. 2014). Wild *C. elegans* isolates used for genome-wide association are described previously (Cook, Zdraljevic, Tanny, et al. 2016; Cook, Zdraljevic, Roberts, et al. 2016). Recombinant inbred advanced intercross lines (RIAILs) used for linkage mapping were constructed previously (Andersen et al. 2015). Strains constructed for this manuscript are listed in Supplemental Information. Construction of individual strains is detailed in the corresponding sections below.

### High-throughput fitness assay

We used a modified version (Supplemental figure 1) of the high-throughput fitness assay (HTA) described previously (Andersen et al. 2015). In short, strains are passaged for four generations to reduce transgenerational effects from starvation or other stresses. Strains are then bleach-synchronized and aliquoted to 96-well microtiter plates at approximately one embryo per microliter in K medium (Boyd et al. 2012). Embryos are then hatched overnight to the L1 larval stage. The following day, hatched L1 animals are fed HB101 bacterial lysate (Pennsylvania State University Shared Fermentation Facility, State College, PA) at a final concentration of 5 mg/ml and grown to the L4 stage after two days at 20°C. Three L4 larvae are then sorted using a large-particle flow cytometer (COPAS BIOSORT, Union Biometrica, Holliston, MA) into microtiter plates that contain HB101 lysate at 10 mg/ml, K medium, 31.25 μM kanamycin, and either drug dissolved in 1% DMSO or 1% DMSO. The animals are then grown for four days at 20°C. During this time, the animals will mature to adulthood and lay embryos that encompass the next generation. Prior to the measurement of fitness parameters from the population, animals are treated with sodium azide (50 mM) to straighten their bodies for more accurate length measurements. Traits that are measured by the BIOSORT include brood size and animal length (time of flight or TOF).

### Calculation of fitness traits for genetic mappings

Phenotype data generated using the BIOSORT were processed using the R package *easysorter*, which was specifically developed for processing this type of data set (Shimko & Andersen 2014). Briefly, the function *read_data*, reads in raw phenotype data, runs a support vector machine to identify and eliminate bubbles. Next, the *remove_contamination* function eliminates any wells that were contaminated prior to scoring population parameters for further analysis. Contamination is assessed by visual inspection. The *sumplate* function is then used to generate summary statistics of the measured parameters for each animal in each well. These summary statistics include the 10th, 25th, 50th, 75th, and 90th quantiles for TOF. Measured brood sizes are normalized by the number of animals that were originally sorted into the well. After summary statistics for each well are calculated, the *regress*(*assay*=*TRUE*) function in the *easysorter* package is used to fit a linear model with the formula (*phenotype ∼ assay*) to account for any differences between assays. Next, outliers are eliminated using the *bamf_prune* function. This function eliminates strain values that are greater than two times the IQR plus the 75th quantile or two times the IQR minus the 25th quantile, unless at least 5% of the strains lie outside this range. Finally, drug-specific effects are calculated using the *regress*(*assay*=*FALSE*) function from *easysorter*, which fits a linear model with the formula (*phenotype ∼ control phenotype*) to account for any differences in population parameters present in control DMSO-only conditions.

### Topoisomerase II poisons dose-response assays

All dose-response experiments were performed on four genetically diverged strains (Bristol, Hawaii, DL238, and JU258) in technical quadruplicates prior to performing GWA and linkage mapping experiments (Supplemental table 4). Animals were assayed using the HTA, and phenotypic analysis was performed as described above. Drug concentrations for GWA and linkage mapping experiments were chosen based on two criteria – an observable drug-specific effect and broad-sense heritability H^2^. We aimed to use the first concentration for which a drug-specific effect with a maximum H^2^ was observed. Broad-sense heritability estimates were calculated using the *lmer* function in the *lme4* package with the following model (*phenotype ∼1* + (*1|strain*)). Concentrations for each chemotherapeutic used in mapping experiments are; etoposide - 250μM, teniposide - 125 μM, amsacrine - 50 μM, dactinomycin - 15 μM, and XK469 - 1000 μM. All topoisomerase II poisons used in this study were purchased from Sigma (XK469 cat#X3628, etoposide cat#E1383, amsacrine cat#A9809, dactinomycin cat#A1410, and teniposide cat#SML0609).

### Linkage Mapping

A total of 265 RIAILs were phenotyped in the HTA described previously for control and etoposide conditions. The phenotype data and genotype data were entered into R and scaled to have a mean of zero and a variance of one for linkage analysis (Supplemental table 5). Quantitative trait loci (QTL) were detected by calculating logarithm of odds (LOD) scores for each marker and each trait as –n(ln(l − R^2^)/2*ln*(10)), where r is the Pearson correlation coefficient between RIAIL genotypes at the marker and phenotype trait values (Bloom et al.2013). The maximum LOD score for each chromosome for each trait was retained from three iterations of linkage mappings (Supplemental table 6). We randomly permuted the phenotype values of each RIAIL while maintaining correlation structure among phenotypes 1000 times to estimate significance empirically. The ratio of expected peaks to observed peaks was calculated to determine the genome-wide error rate of 5% of LOD 4.61. Broad-sense heritability was calculated as the fraction of phenotypic variance explained by strain from fit of a linear mixed-model of repeat phenotypic measures of the parents and RIAILs (Brem & Kruglyak 2005). The total variance explained by each QTL was divided by the broad-sense heritability to determine how much of the heritability is explained by each QTL. Confidence intervals were defined as the regions contained within a 1.5 LOD drop from the maximum LOD score.

### Genome-wide association mapping

Genome-wide association (GWA) mapping was performed using 152 *C. elegans* isotypes (Supplemental table 7). We used the *cegwas* R package for association mapping (Cook, Zdraljevic, Roberts, et al. 2016). This package uses the EMMA algorithm for performing association mapping and correcting for population structure (Kang et al. 2008), which is implemented by the GWAS function in the *rrBLUP* package (Endelman 2011). The kinship matrix used for association mapping was generated using a whole-genome high-quality single-nucleotide variant (SNV) set (Cook, Zdraljevic, Tanny, et al. 2016) and the *A.mat* function from the *rrBLUP* package. SNVs previously identified using RAD-seq (Andersen et al. 2012) that had at least 5% minor allele frequency in the 152 isotype set were used for performing GWA mappings. Association mappings that contained at least one SNV that had a −*log*_10_(*p*) value greater than the Bonferroni-corrected value were processed further using fine mapping (Supplemental table 8). Tajima’s D was calculated using the *tajimas_d* function in the *cegwas* package using default parameters (window size = 300 SNVs, sliding window distance = 100 SNVs, outgroup = N2) (Supplemental table 9).

### Fine mapping

Fine mapping was performed on variants from the whole-genome high-quality SNV set within a defined region of interest for all mappings that contained a significant QTL. Regions of interest surrounding a significant association were determined by simulating a QTL with 20% variance explained at every RAD-seq SNV present in 5% of the phenotyped population. We then identified the most correlated SNV for each mapping. Next, we determined the number of SNVs away from the simulated QTL SNV position that captured 95% of the most correlated SNVs. A range of 50 SNVs upstream or downstream of the peak marker captured 95% of the most significant SNVs in the simulated mappings. We therefore used a region 50 SNVs from the last SNV above the Bonferroni-corrected *p*-value on the left side of the peak marker and 50 SNVs from the last SNV above the Bonferroni-corrected *p*-value on the right side of the peak marker.

The *snpeff* function from the *cegwas* package was used to identify SNVs from the whole-genome SNV set with high to moderate predicted functional effects present in a given region of interest (Cingolani et al. 2012). The correlation between each variant in the region of interest and the kinship-corrected phenotype used in the GWA mapping was calculated using the *variant_correlation* function and processed using the *process_correlations* function in the *cegwas* package (Supplemental table 10). ClustalX was used to perform the multiple sequence alignment between various topoisomerase II orthologs (Supplemental file 1).

### Near-isogenic line generation

NILs were generated by crossing N2xCB4856 RIAILs to each parental genotype. For each NIL, eight crosses were performed followed by six generations of selfing to homozygose the genome. Reagents used to generate NILs are detailed in Supplemental Information. The NILs responses to 250 μM etoposide were quantified using the HTA fitness assay described above (Supplemental table 11).

### Dominance tests

Dominance experiments were performed using the fluorescent reporter strain EG7952 *oxTi207 [eft-3p*::*GFP::unc-54 3’UTR* + *hsp::peel-1* + *NeoR* + *Cbr-unc-119*(+)]. Hermaphrodites of N2 and CB4856 were crossed to male EG7952 reporter strain, which expresses GFP, to ensure that we could measure heterozygous cross progeny by the presence of GFP. Three GFP-positive progeny were manually transferred to a 96-well assay microtiter plate containing to 250 μM etoposide dissolved in 1% DMSO or 1% DMSO control, in addition to K medium, HB101 lysate at 10 mg/ml, and 31.25 μM kanamycin. Animals were grown for four days at 20°C. The phenotypes of the progeny were scored using the BIOSORT as described above (Supplemental table 12). Heterozygous progeny were computationally identified as those individuals that had fluorescence levels between the non-fluorescent and fluorescent parental strains.

### *top-2* and *npp-3* Complementation

To perform the complementation experiments, N2 and CB4856 males were both crossed to both VC1474 *top-2*(*ok1930*)*/mIn1 [mIs14 dpy-10*(*e128*)] and VC1505 *npp-3*(*ok1900*)*/mIn1 [mIs14 dpy-10*(*e128*)] hermaphrodites. Three non-GFP L4 hermaphrodite progeny were manually picked into experimental wells containing either 250 μM etoposide dissolved in 1% DMSO or 1% DMSO without etoposide, in addition to HB101 lysate at 10 mg/ml, K medium, and 31.25 μM kanamycin. Animals were grown for four days at 20°C. The phenotypes of the progeny were scored using the BIOSORT as described above (Supplemental table 13).

### *top-2* reciprocal hemizygosity

VC1474 *top-2*(*ok1930*)*/mIn1 [mIs14 dpy-10*(*e128*)] was used for *top-2* complementation tests. *top-2*(*ok1930*) and *mIn1[mIs14 dpy-10*(*e128*)] were individually introgressed into CB4856 for 10 generations. Once individual crosses were completed, CB4856 *mIn1 [mIs14 dpy-10(e128)]* was crossed to CB4856 *top-2*(*ok1930*) to generate ECA338, which contains a *mIn1-*balanced *top-2*(*ok1930*). oECA1003 and oECA1004 were used to verify the presence of *top-2*(*ok1930*) during crosses.

To perform the reciprocal hemizygosity experiment, N2 and CB4856 males were both crossed to both VC1474 and ECA338 hermaphrodites. Three non-GFP L4 hermaphrodite progeny were manually picked into experimental wells containing either 250 μM etoposide dissolved in 1% DMSO or 1% DMSO without etoposide, in addition to HB101 lysate at 10 mg/ml, K medium, and 31.25 μM kanamycin. Animals were grown for four days at 20°C. The phenotypes of the progeny were scored using the BIOSORT as described above (Supplemental table 14).

### Generation of *top-2* allele replacement strains

All allele replacement strains were generated using CRISPR/Cas9-mediated genome engineering, using the co-CRISPR approach (Kim et al. 2014) with Cas9 ribonucleoprotein delivery (Paix et al. 2015). Alt-R™ crRNA and tracrRNA (Supplemental Information) were purchased from IDT (Skokie, IL). tracrRNA (IDT, 1072532) was injected at a concentration of 13.6 μM. The *dpy-10* and the *top-2* crRNAs were injected at 4 μM and 9.6 μM, respectively. The *dpy-10* and the *top-2* single-stranded oligodeoxynucleotides (ssODN) repair templates were injected at 1.34 μM and 4 μM, respectively. Cas9 protein (IDT, 1074182) was injected at 23 uM. To generate injection mixes, the tracrRNA and crRNAs were incubated at 95°C for 5 minutes and 10°C for 10 minutes. Next, Cas9 protein was added and incubated for 5 minutes at room temperature. Finally, repair templates and nuclease-free water were added to the mixtures and loaded into pulled injection needles (1B100F-4, World Precision Instruments, Sarasota, FL). Individual injected *P_0_* animals were transferred to new 6 cm NGM plates approximately 18 hours after injections. Individual *F_1_* rollers were then transferred to new 6 cm plates and allowed to generate progeny. The region surrounding the desired Q797M (or M797Q) edit was then amplified from *F_1_* rollers using oECA1087 and oECA1124. The PCR products were digested using the *Hpy*CH4III restriction enzyme (R0618L, New England Biolabs, Ipswich, MA). Differential band patterns signified successfully edited strains because the N2 Q797, which is encoded by the CAG codon, creates an additional *Hpy*CH4III cut site. Non-Dpy, non-Rol progeny from homozygous edited *F_1_* animals were propagated. If no homozygous edits were obtained, heterozygous *F_1_* progeny were propagated and screened for presence of the homozygous edits. *F_1_* and *F_2_* progeny were then Sanger sequenced to verify the presence of the proper edit. Allele swap strains responses to the topoisomerase II poisons were quantified using the HTA fitness ass described above.

### Molecular docking simulations

The *C. elegans* TOP-2 three-dimensional structure homology model was built by threading the *C. elegans* TOP-2 peptide to the human topoisomerase II beta structure (PDB accession code 3QX3; 59% identity, 77% similarity) using the Prime3.1 module implemented in Schrodinger software (Jacobson et al. 2002; Jacobson et al. 2004). After building the model, a robust energy minimization was carried out in the Optimized Potentials for Liquid Simulations (OPLS) force field. The minimized structure was subjected to MolProbity analysis, and the MolProbity score suggested with greater than 95% confidence that the minimized structure model was a good high-resolution structure (Davis et al. 2007).

Next, the Prot-Prep wizard was used to prepare the TOP-2 homology model, which fixed the hydrogen in the hydrogen bond orientations, eliminated the irrelevant torsions, fixed the missing atoms, assigned the appropriate force field charges to the atoms (Sastry et al. 2013). After preparing the structure, the glutamine 797 was mutated to various rotamers of methionine (Q797M), which subsequently underwent minimization in the OPLS force field. The energy-minimized structure was used in the *in silico* experiments.

The structure data file of etoposide (DrugBank ID: DB00773) was obtained from PubMed and was subjected to ligand preparation panel of the Schrodinger software. Using the induced fit docking (IFD) module of Schrodinger and Suflex software, we carried out the docking of etoposide with the glutamine (Supplemental file 2A) and methionine (Supplemental file 2B) forms of the *C. elegans* TOP-2 homology model. After the docking experiments, we analyzed the docked poses of the ligands bound to the TOP-2 homology models from both docking engines. Change in free energy (ΔG) and the hydrophobicity parameter were calculated using Schrodinger.

### CRISPR-Cas9 gene editing in human cells

Gene-editing experiments were performed in human 293T cells (ATCC) grown in DMEM with 10% FBS. On day zero, 500,000 cells were seeded per well in a six-well plate format. The following day, two master mixes were prepared: a) LT-1 transfection reagent (Mirus) was diluted 1:10 in Opti-MEM and incubated for five minutes; b) a DNA mix of 500 ng Cas9-sgRNA plasmid with 250 pmol repair template oligonucleotide (Supplemental Information) was diluted in Opti-MEM in a final volume of 100 μL. 100 μL of the lipid mix was added to each of the DNA mixes and incubated at room temperature for 25 minutes. Following incubation, the full 200 μL volume of DNA and lipid mix was added drop-wise to the cells, and the cells were centrifuged at 1000xg for 30 min. Six hours post-transfection, the media on the cells was replaced, and the cells were passaged as needed. On day six, five million cells from each condition were pelleted to serve as an early time point for the editing efficiency, and five million cells were then passaged on the five drugs at two doses for 12 days, at which time all surviving cells were pelleted. Concentrations used for each small molecule are: etoposide – 500 nM, 100 nM; amasacrine – 500 nM, 100nM; teniposide – 20 nM, 4 nM; dactinomycin – 4 nM, 800 pM; and XK469 – 5 μM, 1μM.

### Analysis of CRISPR-Cas9 editing in human cells

gDNA was extracted from cell pellets using the QIAGEN (QIAGEN, Hilden, Germany) Midi or Mini Kits based on the size of the cell pellet (cat # 51183, 51104) according to the manufacturer’s recommendations. TOP2A and B loci were first amplified with 17 cycles of PCR using a touchdown protocol and the NEBnext 2x master mix (New England Biolabs M0541). The resulting product served as input to a second PCR, using primers that appended a sample-specific barcode and the necessary adaptors for Illumina sequencing. The resulting DNA was pooled, purified with SPRI beads (A63880, Beckman Coulter, Brea, CA), and sequenced on an Illumina MiSeq with a 300 nucleotide single-end read with an eight nucleotide index read. For each sample, the number of reads exactly matching the wild-type and edited TOP2A/B sequence were determined (Supplemental table 15).

## Acknowledgments

The authors would like to thank Samuel Rosenberg for assistance on early mappings of drug sensitivities, Rama Mishra of the Center for Molecular Innovation and Drug Discovery core for molecular dynamic simulations, Mudra Hegde of the Broad Institute for assistance with sequence analysis, and members of the Andersen laboratory for critical reading of this manuscript, and Joshua Bloom for editorial comments. This work was supported by a National Institutes of Health R01 subcontract to E.C.A. (GM107227), the Chicago Biomedical Consortium with support from the Searle Funds at the Chicago Community Trust, a Sherman-Fairchild Cancer Innovation Award to E.C.A., and an American Cancer Society Research Scholar Grant to E.C.A. (127313-RSG-15-135-01-DD), along with support from the Cell and Molecular Basis of Disease training grant (T32GM008061) to S.Z., from the Life Science Research Foundation award to H.S.S., and from the National Science Foundation Graduate Research Fellowship (DGE-1324585) to D.E.C. J.G.D. is a Merkin Institute Fellow and is supported by the Next Generation Fund at the Broad Institute of MIT and Harvard.

---

**Supplemental Figure 1.**
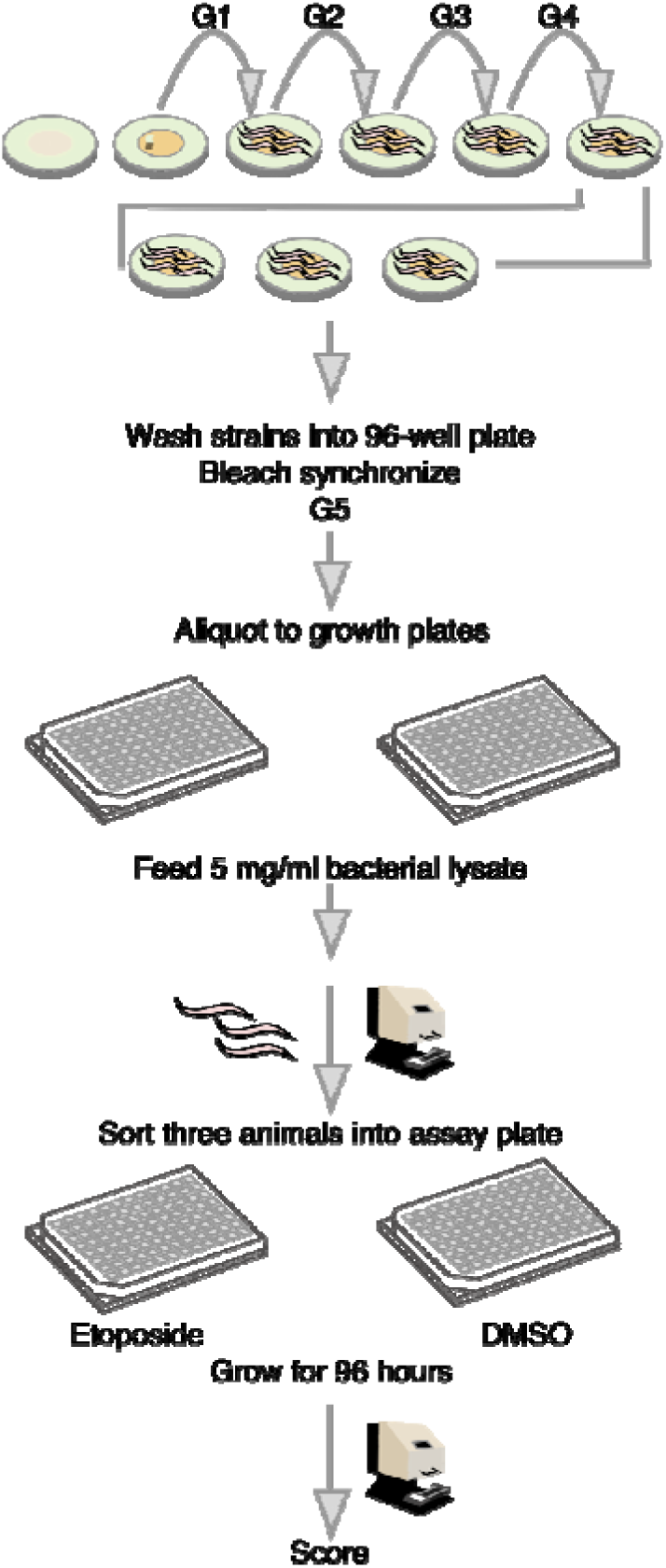
A schematic of the high-throughput phenotyping assay is shown. Individual strains are passaged for four generations on agar plates seeded with OP50 bacteria by transfer of seven L4 larval stage animals to a fresh plate each generation every three days. Animals are bleach synchronized and aliquoted to 96-well microtiter plates in 50 μl of K medium at a concentration of one embryo per μL. Aliquoted embryos are incubated overnight at 20°C. The following day, 5 μl of 50 mg/ml HB101 lysate is added to each well. Animals are then grown for two days to the L4 larval stage. Then, three L4 animals are sorted to assay plates containing drug or DMSO using the BIOSORT. Four days later, 200 μl M9 plus 50 mM sodium azide is added to each well and strains are scored using the BIOSORT.

**Supplemental Figure 2.**
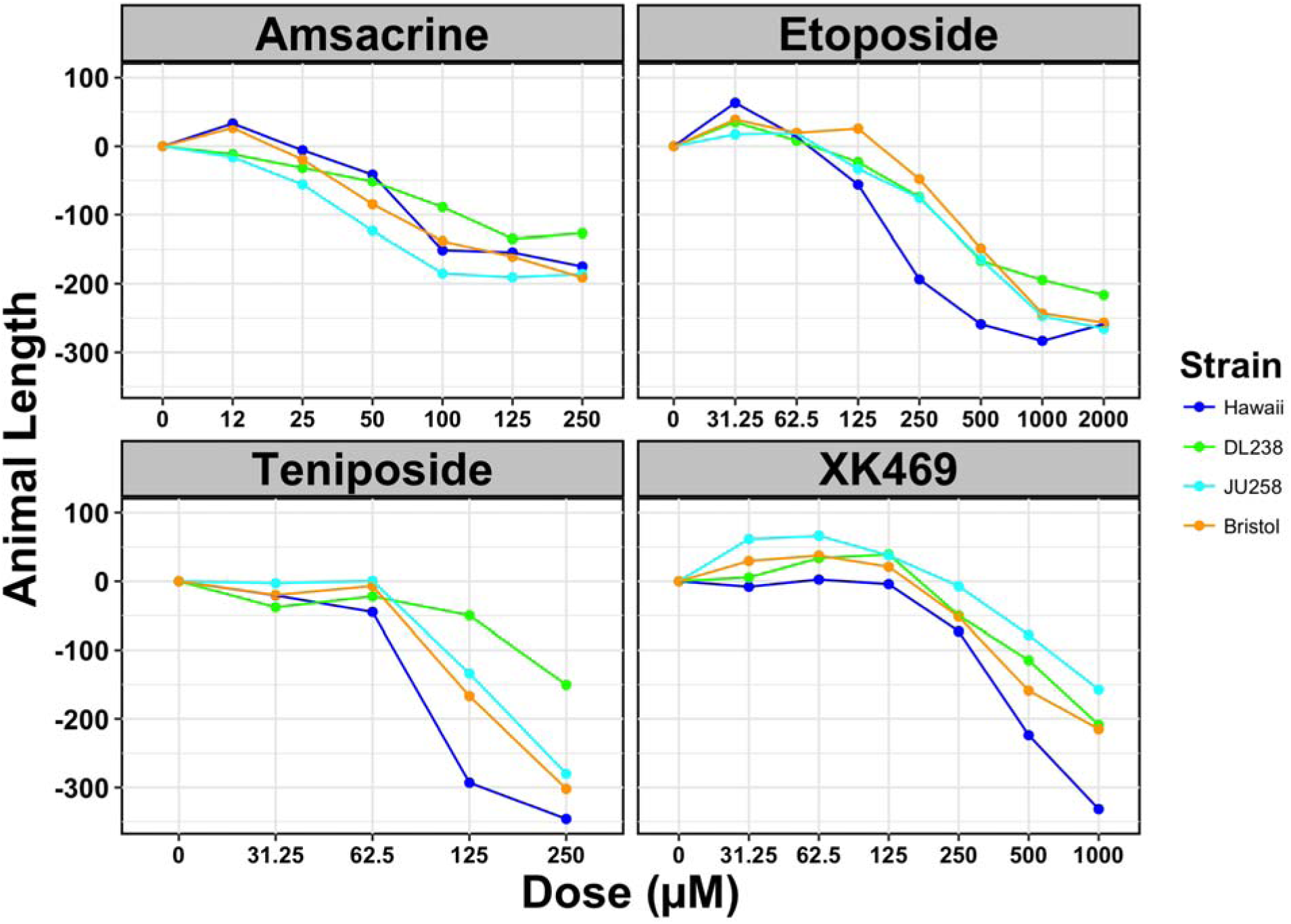
Dose response curves of four genetically diverged *C. elegans* strains exposed to the topoisomerase II poisons used in this study are shown. For each drug, concentration in μΜ is plotted on the x-axis, and the regressed 75th quantile of animal length is plotted on the y-axis. The red numbers above the dose response lines correspond to broad-sense heritability estimates. The heritability estimates for the drug concentrations used in experiments throughout this study are: *H^2^* = 0.47 for amsacrine at 50 μΜ, *H^2^* = 0.62 for etoposide at 250 μΜ, *H^2^* = 0.73 for teniposide at 125 μΜ, and *H^2^* = 0.9 for XK469 at 1000 μΜ. The Bristol strain (N2) is colored in orange, the Hawaiian strain (CB4856) in blue, JU258 in cyan, and DL238 in green.

**Supplemental Figure 3.**
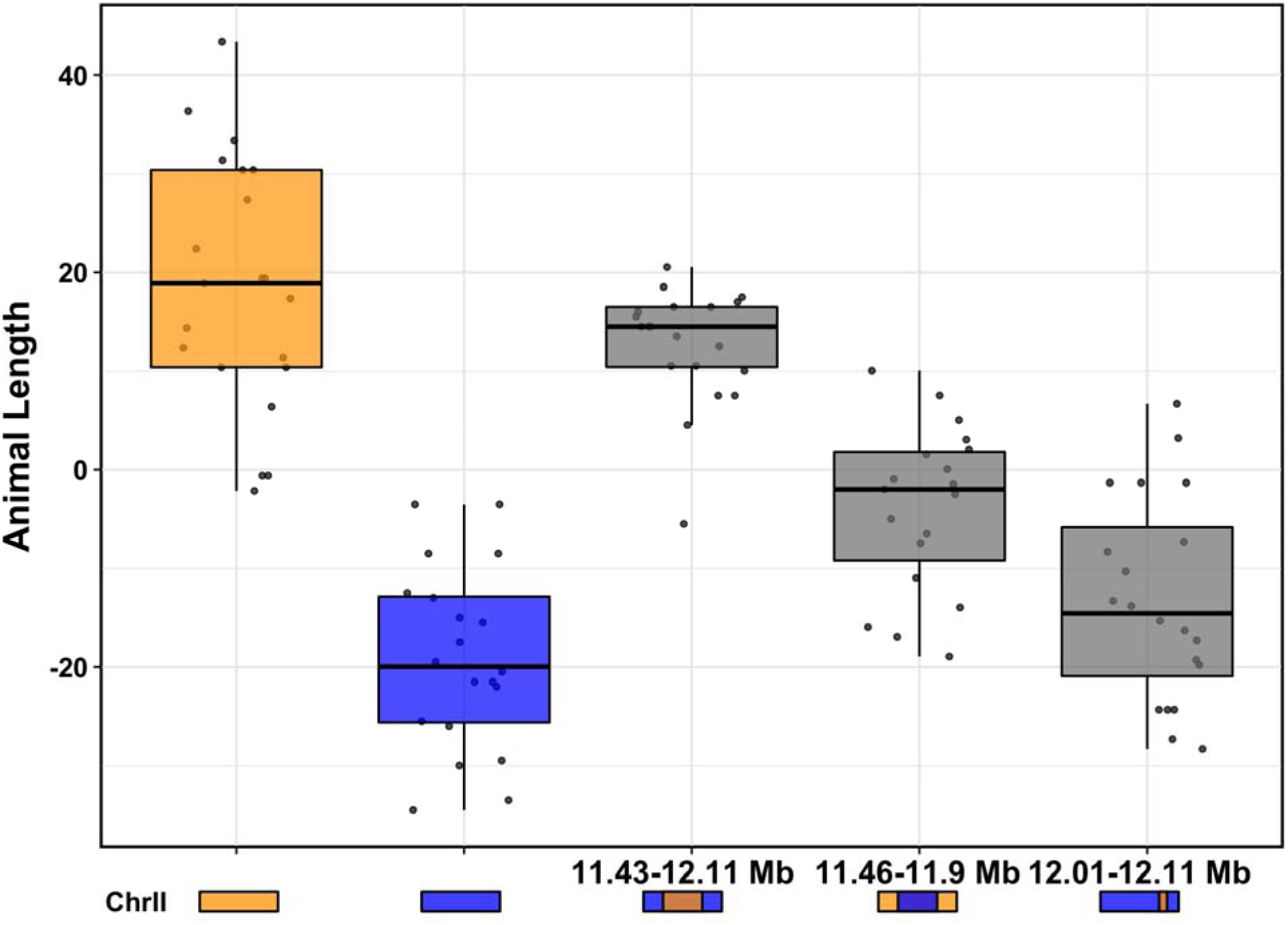
Tukey box plots of NIL regressed median animal length in the presence of etoposide are shown. NIL genotypes are indicated below the plot by colored rectangles, Bristol (orange) or Hawaii (blue). Numbers above rectangles correspond to the genomic position of the introgression region on chromosome II.

**Supplemental Figure 4.**
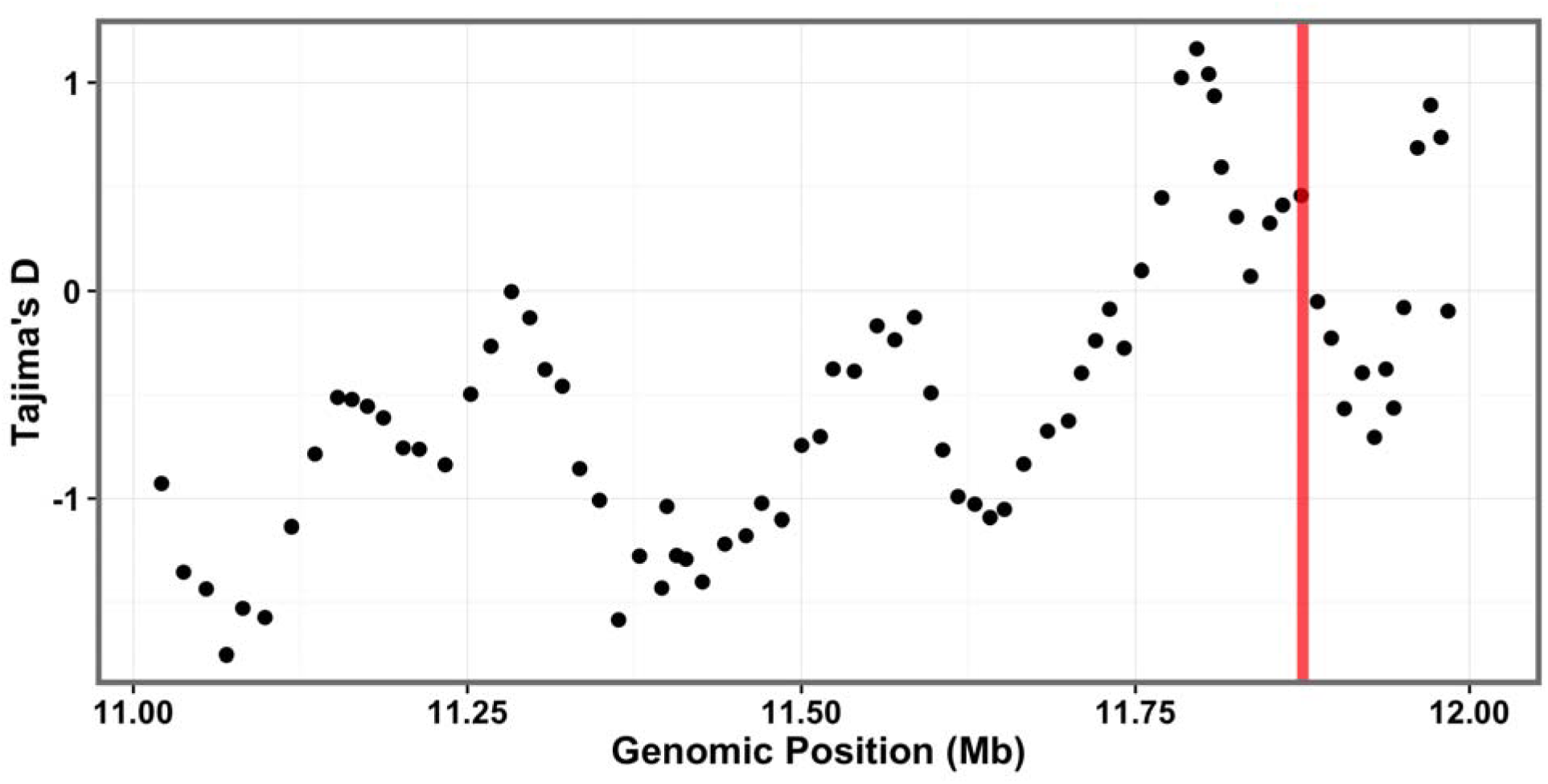
Divergence, as measured by Tajima’s D, is shown across the etoposide QTL confidence interval (II:11021073-12008179). The whole-genome SNV data set (Cook, Zdraljevic, Tanny, et al. 2016; Cook, Zdraljevic, Roberts, et al. 2016) was used for Tajima’s D calculations. Window size for the calculations was 300 SNVs with a 100 SNV sliding window size. The vertical red line marks the position of the *top-2* locus. The *tajimas_d* function in the *cegwas* package was used to perform the calculations.

**Supplemental Figure 5.**
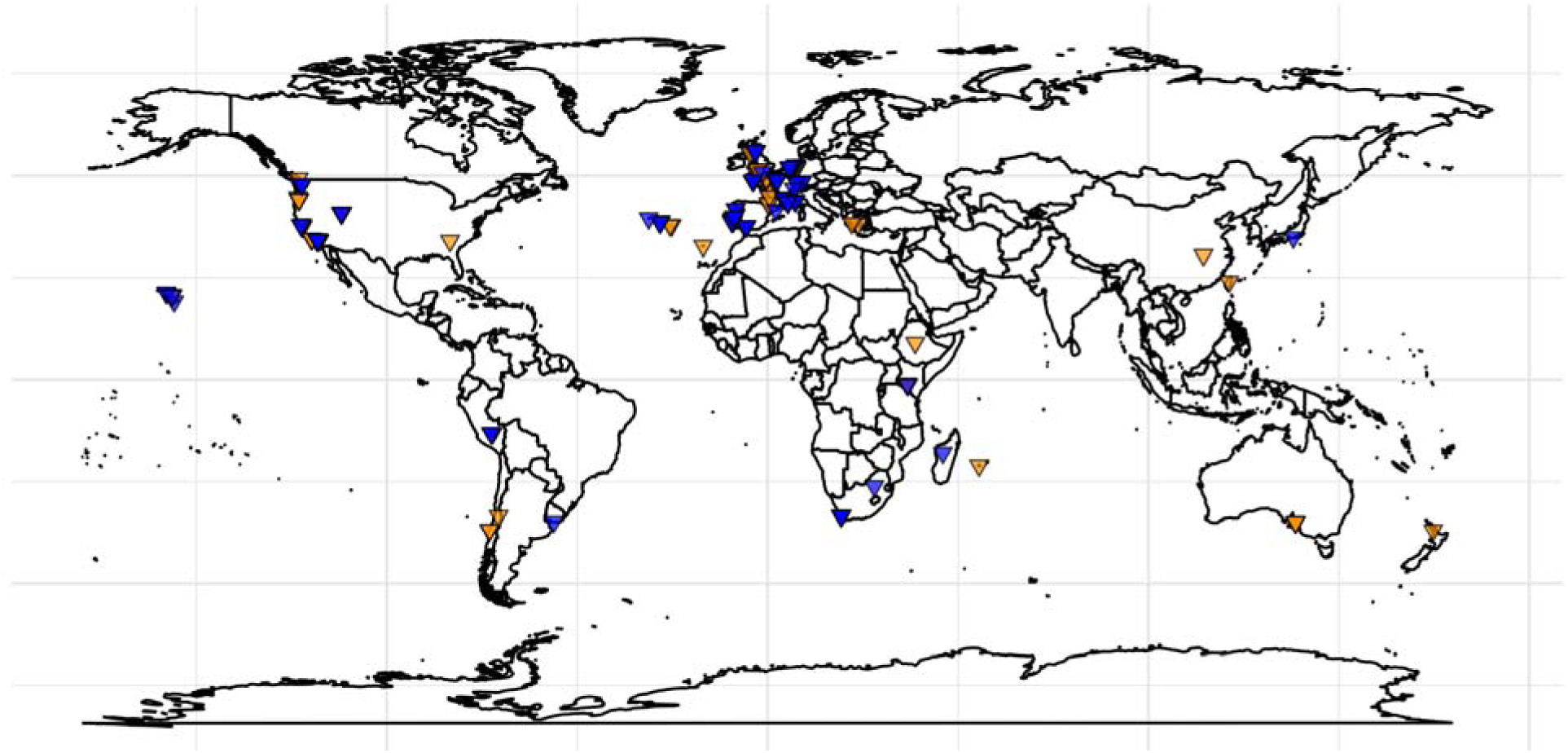
The worldwide distribution of the TOP-2(Q797M) allele is shown. Glutamine (REF) is shown in orange and methionine (ALT) is shown in blue. Latitude and longitude coordinates of sampling locations were used to plot individual strains on the map (Cook, Zdraljevic, Roberts, et al. 2016).

**Supplemental Figure 6.**
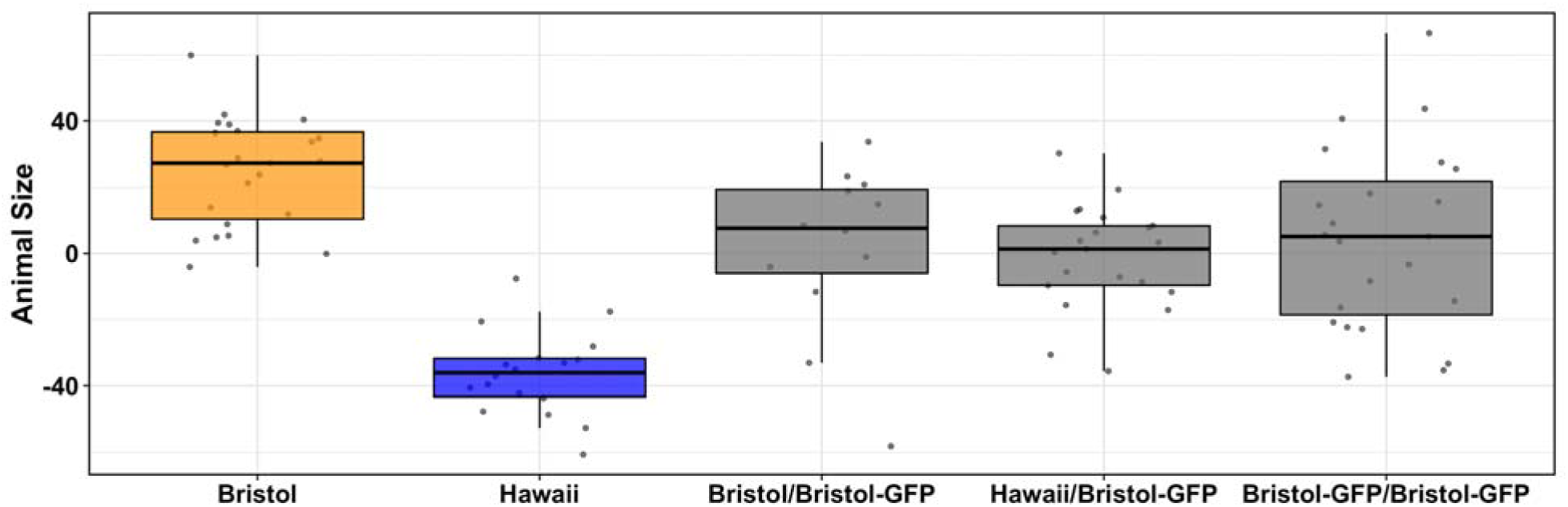
Etoposide resistance is dominant. Tukey box plots of Bristol and Hawaii regressed median animal length in response to etoposide are plotted in orange and blue, respectively. GFP-containing Bristol strains (EG7952) were crossed to the Bristol and Hawaii strains and heterozygous progeny were assayed using the high-throughput fitness assay. Tukey box plots of heterozygotes are shown in gray. All heterozygote strains are significantly different from the parental Bristol and Hawaii strains (Tukey HSD, *p* < 0.02). All heterozygote strains are not significantly different from each other (Tukey HSD, *p* > 0.91).

**Supplemental Figure 7.**
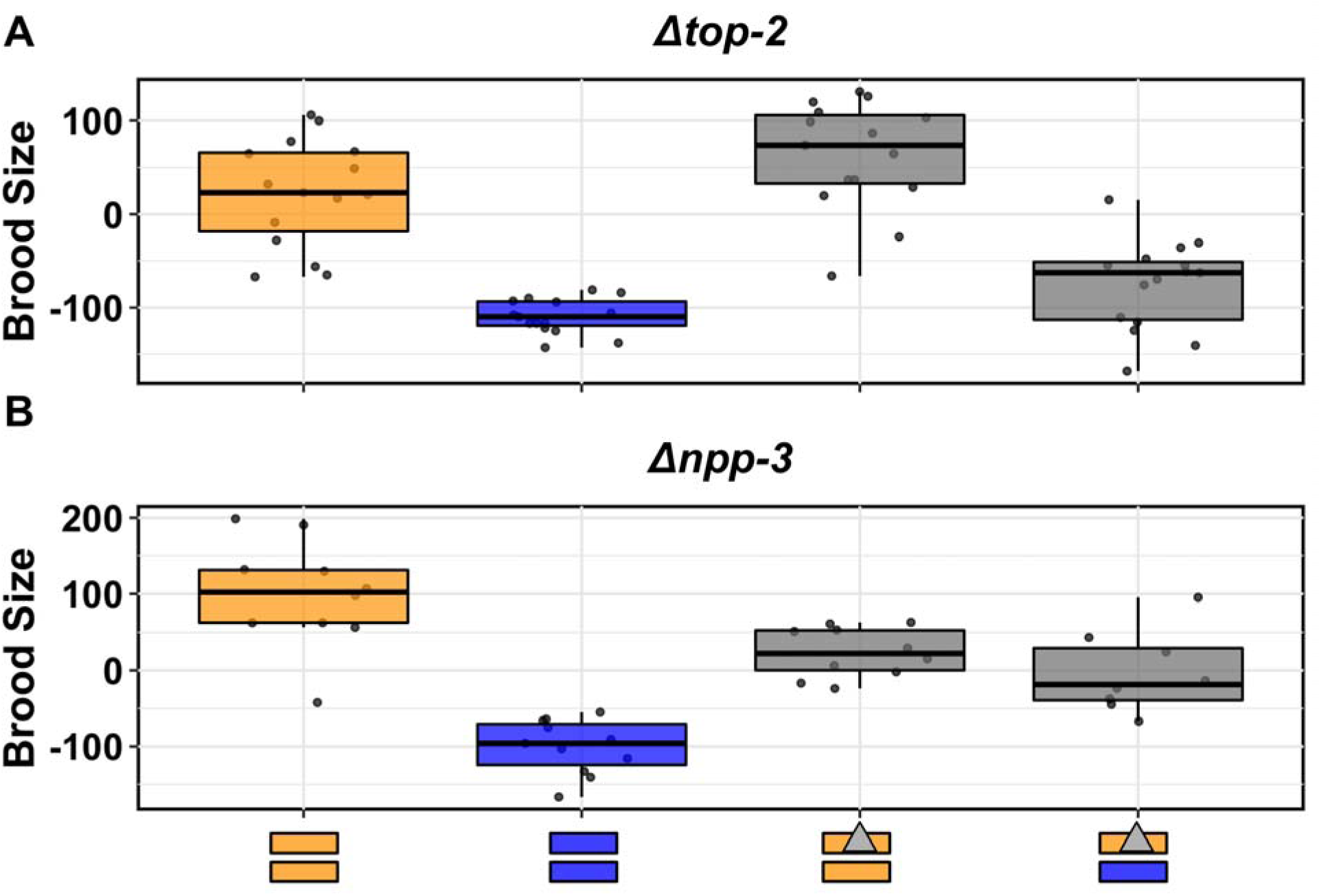
Complementation tests of the *npp-3* and *top-2* genes are shown. Tukey box plots of the residual brood size distribution of Bristol (orange) and Hawaii (blue) compared to two heterozygous (A) *top-2* and (B) *npp-3* deletion strains (gray) in response to etoposide are plotted. Orange (Bristol) and blue (Hawaii) rectangles below the plot correspond to the two chromosome II homolog genotypes. Gray triangles denote chromosomes with the *top-2* deletion allele. (A) The Bristol strain and the Bristol/Bristol(Δ*top-2*) heterozygous strain are not significantly different from each other (Tukey HSD, *p*-value 0.104). The Hawaii strain and the Hawaii/Bristol(Δ*top-2*) heterozygous strain are not significantly different from each other (Tukey HSD, *p*-value 0.226). All other comparisons are significant (Tukey HSD *p*-value < 4.1E-6). (B) The Hawaii/Bristol(Δ*npp-3*) and the Bristol/Bristol(Δ*npp-3*) heterozygous strain are not significantly different from each other (Tukey HSD, *p*-value 0.68), but all other comparisons are significant (Tukey HSD *p*-value < 0.009).

**Supplemental Figure 8.**
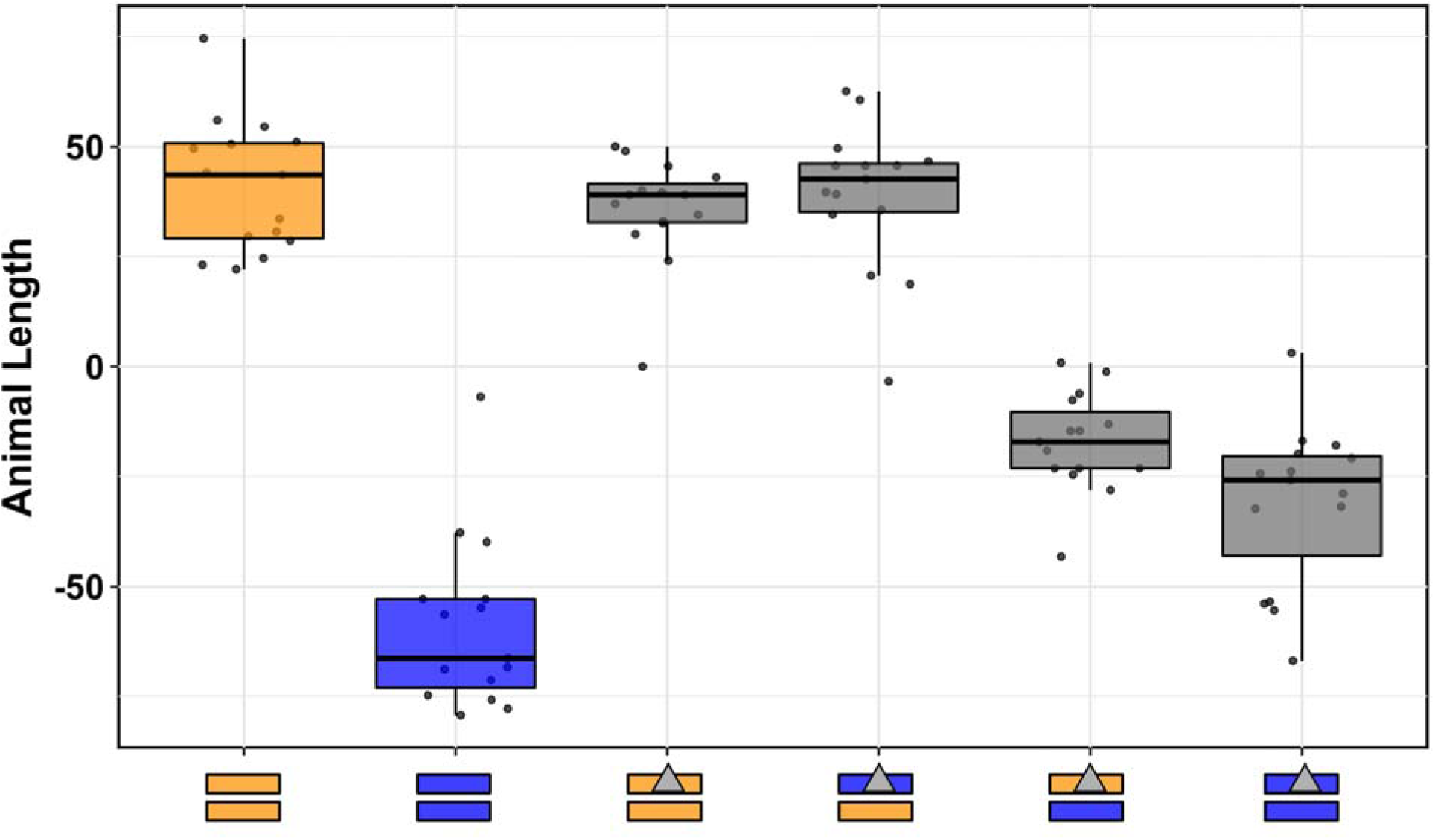
Reciprocal hemizygosity of *top-2* is shown. (A) Tukey box plots of the residual median animal length distribution of the Bristol (orange) and Hawaii (blue) strains compared to four heterozygous *top-2* deletion strains (gray) in response to etoposide are plotted. Orange (Bristol) and blue (Hawaii) rectangles below the plot correspond to the two chromosome II homolog genotypes. Gray triangles denote chromosomes with the *top-2* deletion allele. Phenotypes of heterozygous deletion strains with the Bristol TOP-2 allele are not significantly different from the parental Bristol strain. The Hawaii/Bristol(Δ*top-2*) and the Hawaii/Hawaii(Δ*top-2*) strain phenotypes are not significantly different from each other (Tukey HSD, *p*-value = 0.16), but are both significantly different from the parental Hawaii strain (Tukey HSD, *p*-value < 1E-7 and *p*-value = 0.0001, respectively).

**Supplemental Figure 9.**
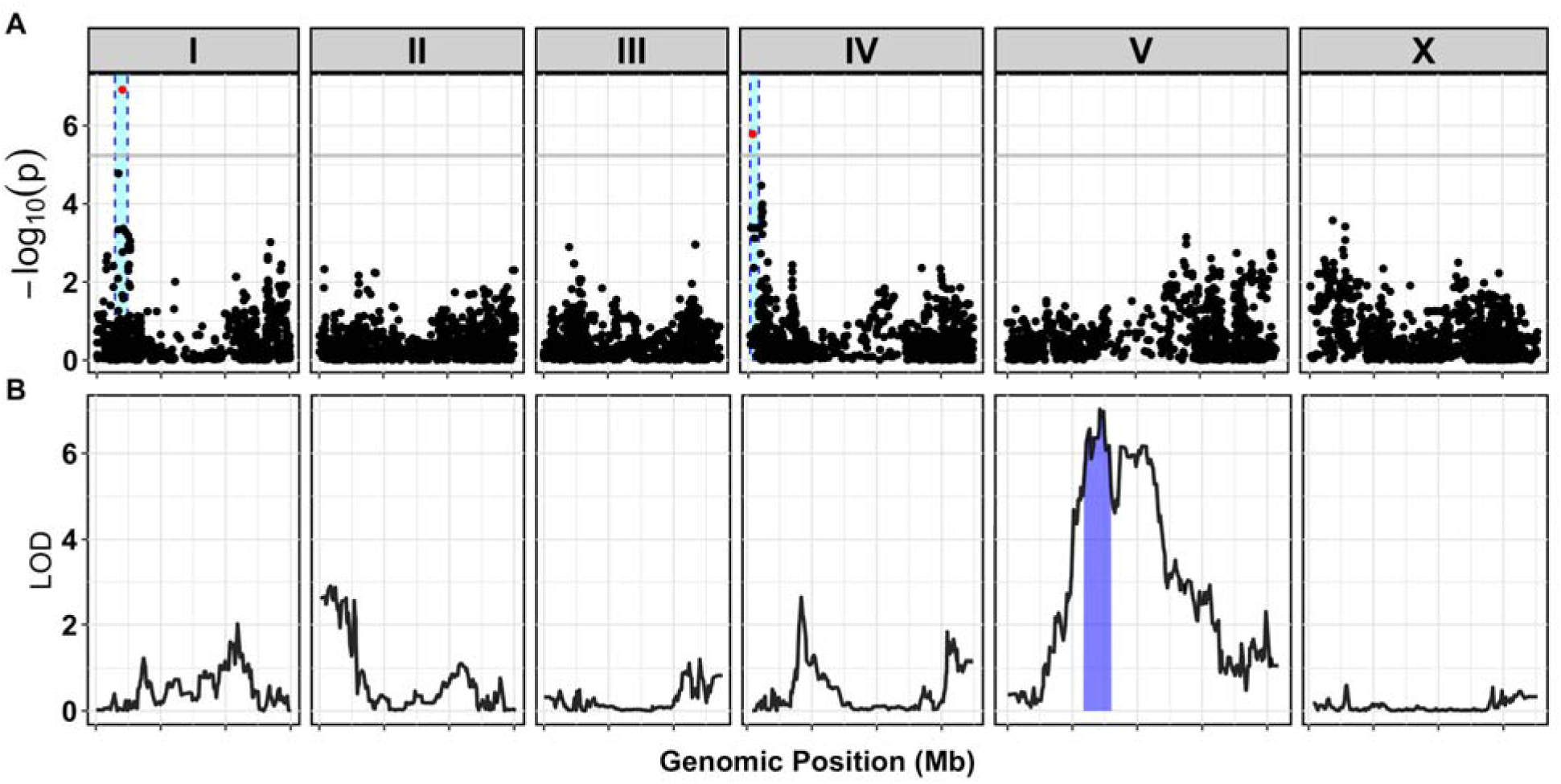
GWA and linkage mapping amsacrine response are shown. (A) A manhattan plot for regressed fraction of animals in the L1 larval stage in the presence of amsacrine is plotted. Each dot represents an SNV that is present in at least 5% of the phenotyped population. The *−log*_10_(*p*) for each SNV is plotted on the y-axis and the genomic position (Mb) is on the x-axis. Each tick on the x-axis corresponds to 5 Mb. SNVs are colored red if they pass the genome-wide Bonferroni-corrected significance threshold, which is denoted by the gray horizontal line. The genomic regions of interest are represented by cyan rectangles surrounding each QTL. (B) A linkage mapping plot for regressed fraction of animals in the L1 larval stage in the presence of amsacrine is shown. The significance value (logarithm of odds, LOD, ratio) is plotted on the y-axis and genomic position (Mb), separated by chromosome, on the x-axis. Each tick on the x-axis corresponds to 5 Mb. The associated 1.5 LOD-drop confidence intervals are represented by blue bars.

**Supplemental Figure 10.**
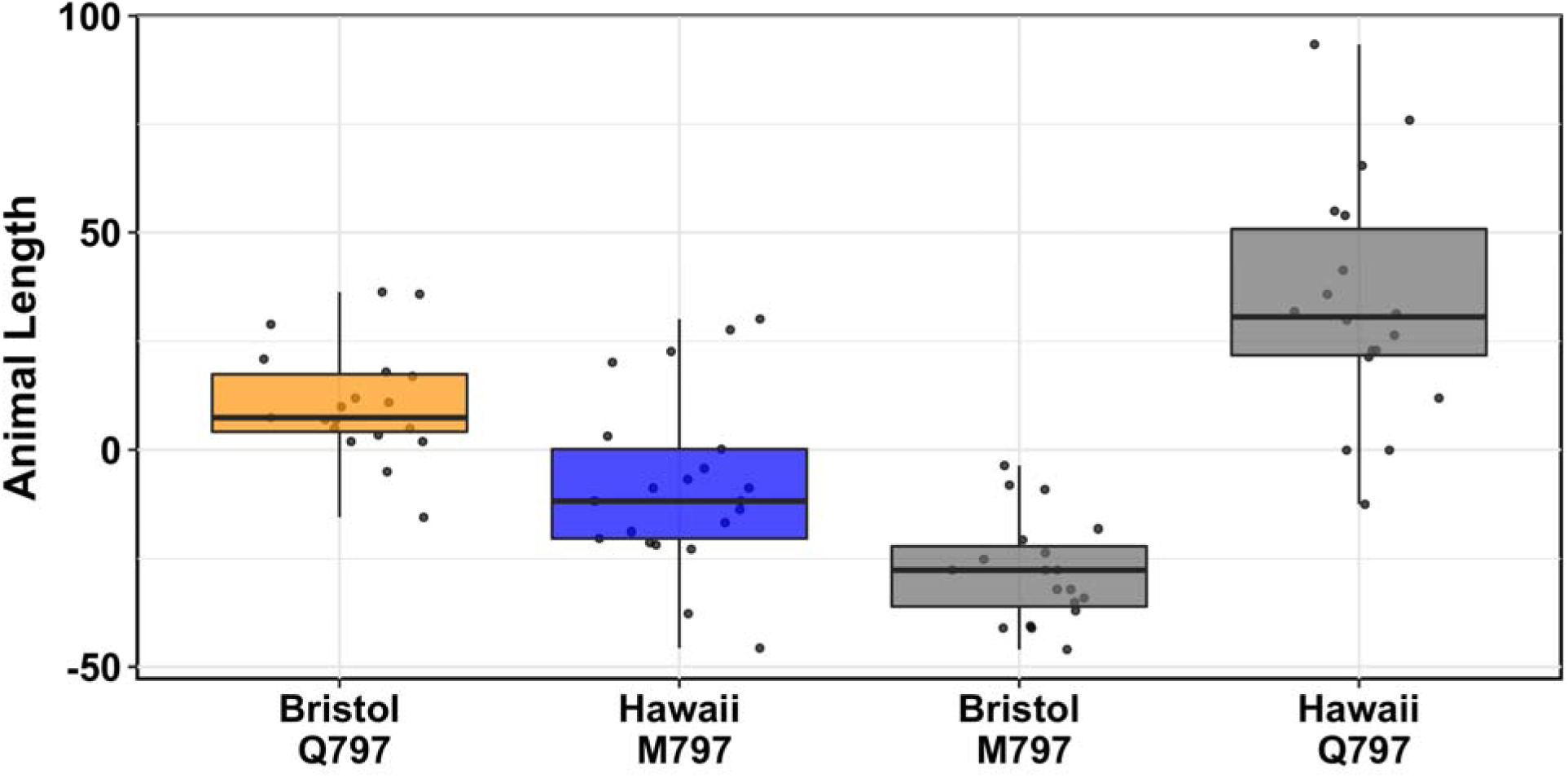
Phenotypic responses to XK469 of allele-replacement strains are shown. Tukey box plots of regressed median animal length in response to XK469 are plotted. Orange corresponds to the Bristol genetic background and blue to Hawaii. Allele-replacement strains are denoted by their genetic background and the corresponding residue change at position 797 of TOP-2. Strains containing the glutamine TOP-2 allele are more resistant to XK469 than strains with the methionine allele. All strain comparisons are significantly different from each other (Tukey HSD, *p*-value < 0.01)

**Supplemental Figure 11.**
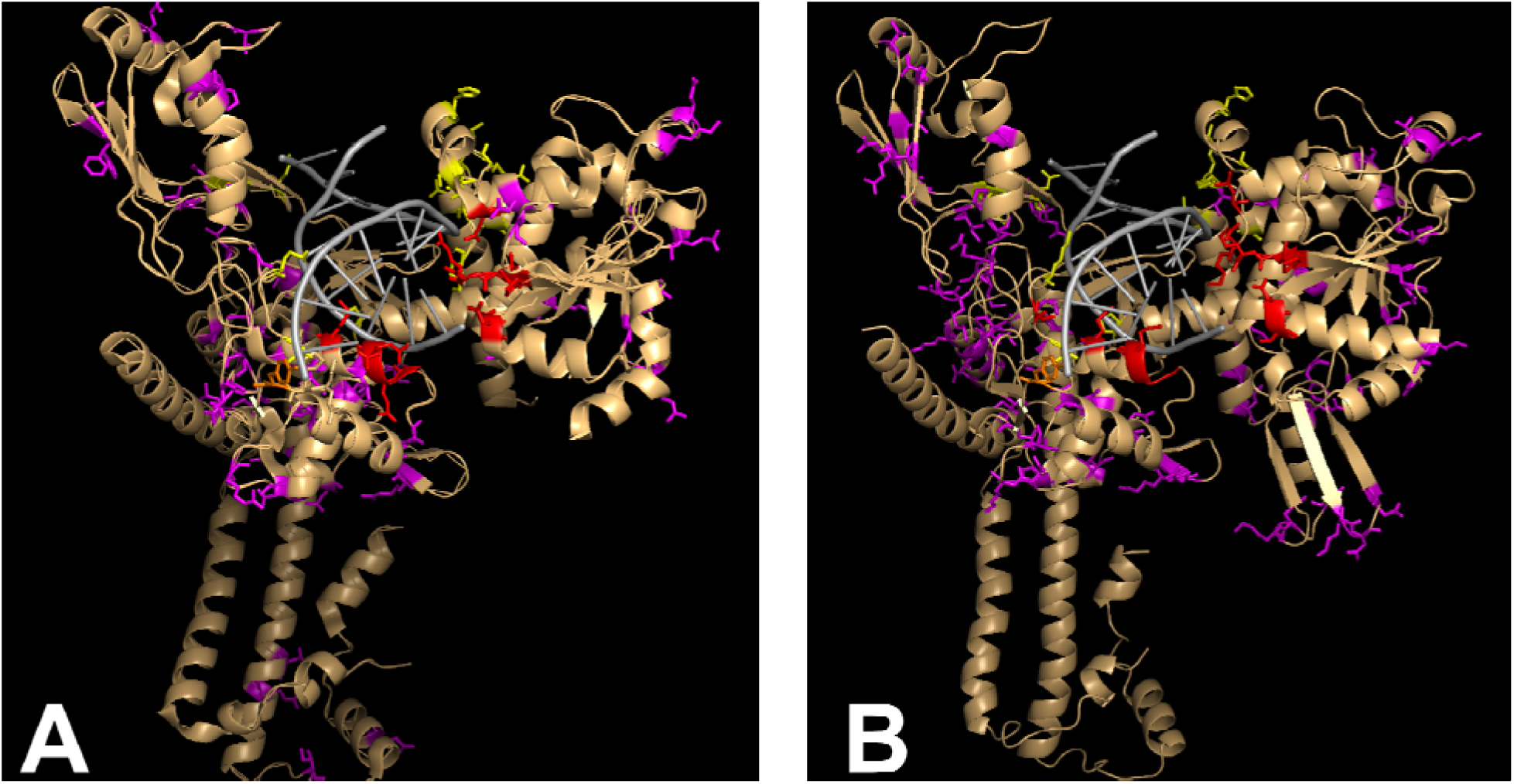
Variant residues in human topoisomerase II enzyme isoforms (A) hTOPOIIα (PDB 4FM9) and (B) hTOPIIβ (PDB 3QX3) are shown. Putative etoposide-binding residues are highlighted in red, DNA-binding residues are highlighted in yellow, and the catalytic tyrosine is highlighted in orange. The cartoon representation of the peptide backbones are shown in tan, and DNA is shown in gray. Residues that vary in the human population are highlighted in purple.

## Supplemental Tables

**Table S1**: CRISPR allele-swap strains phenotype data: Processed phenotype data of CRISPR allele-swap strains in the presence of etoposide.

**Table S2**: hTOP2A variants: Variants present in hTOP2A.

**Table S3**: hTOP2B variants: Variants present in hTOP2B.

**Table S4**: Dose response phenotype data: Processed dose-response phenotype data used for establishing the doses to use for mapping experiments.

**Table S5**: RIAIL phenotype data: Processed phenotype data of RIAILs in the presence of amsacrine or etoposide used to perform linkage mapping.

**Table S6**: Linkage mapping LOD scores: Processed linkage mapping LOD scores at markers across the genome for median animal length in the presence of etoposide.

**Table S7**: Wild isolate phenotype data: Processed phenotype data of wild isolates in the presence of amsacrine or etoposide used to perform GWA mapping.

**Table S8**: GWA mapping significance values: Processed GWA mapping significance values for markers across the genome in the presence of amsacrine of etoposide.

**Table S9**: Tajima’s D values: Sliding window Tajimas’s D values for the etoposide median animal length confidence interval.

**Table S10**: Variant gene correlation values: Spearman’s *rho* values for all coding variants present in the etoposide median animal length confidence interval.

**Table S11**: NIL phenotype data: Processed median animal length phenotype data of NILs in the presence of etoposide.

**Table S12**: Dominance test phenotype data: Processed median animal length phenotype data of strains used to perform the dominance test in the presence of etoposide.

**Table S13**: *npp-3* and *top-2* complementation test phenotype data: Processed brood size phenotype data of strains used to perform the *npp-3* and *top-2* complementation tests in the presence of etoposide.

**Table S14**: *top-2* reciprocal hemizygosity test phenotype data: Processed median animal length phenotype data of strains used to perform the *top-2* reciprocal hemizygosity test in the presence of etoposide.

**Table S15**: hTOP2A and hTOP2B read counts: Processed read counts of hTOP2A and hTOP2B prior to and after exposure to amsacrine, dactinomycin, etoposide, teniposide, XK469, or no drug.

## Supplemental Files

**File S1**: TOP2 multiple sequence alignment: Multiple sequence alignment of multiple TOP2 orthologues.

**File S2**: *C. elegans* TOP-2(Q797) homology model: *C. elegans* TOP-2(Q797) 3-dimensional homology model.

**File S3**: *C. elegans* TOP-2(M797) homology model: *C. elegans* TOP-2(M797) 3-dimensional homology model.

## Supplemental Methods

### Generation of NIL strains

RIAILs that switched genotypes near indel markers were used as starting strains to generate all NILs used in this study. The RIAIL QX327 was used to generate ECA219 *eanIR139[II, CB*>*N2]*,QX322 was used to generate ECA216 *eanIR136[II, N2*>*CB4856]*, and QX327 was used to generate ECA220 *eanIR140[II, N2*>*CB4856]*. Primers oECA605, oECA606, oECA607, and oECA608 were used to follow two indels at 11.64 and 11.91Mb on chromosome II to generate the Hawaii NIL ECA219. Primers oECA593, oECA594, oECA595, and oECA596 were used to follow two indels at 11.43 and 12.11Mb on chromosome II to generate the Bristol NIL ECA216. Primers oECA601, oECA602, oECA03, and oECA604 were used to follow two indels at 12.01 and 12.1Mb on chromosome II to generate the Bristol NIL ECA220. For each NIL, eight crosses were performed followed by six generations of selfing to homozygose the genome. All PCRs were multiplexed and performed with four primers at an annealing temperature of 58°C for 34 cycles with Taq polymerase.

### Generation of Hawaii *Δtop-2* strain

Heterozygous Bristol *top-2*(*ok1930*) and Bristol *mIn1* strains were then crossed to the Hawaiian strain. Males from these crosses were used to outcross to CB4856 for 10 consecutive generations. The primers oECA1003 and oECA1004 were used to follow the *top-2*(*ok1930*)allele during the back crosses. *mIn1* crosses were followed by identifying green fluorescent male progeny. At the end of 10 generations, the CB4856 crosses to *top-2*(*ok1930*) and *mIn1* were crossed to each other to generate a *top-2*(*ok1930*)*/mIn1* strain in the CB4856 genetic background and named ECA338.

### Generation of top-2 allele replacement strains

All allele replacement strains were generated using CRISPR/Cas9-mediated genome engineering, using the co-CRISPR approach described in the main text methods (Kim *et al*. 2014) with Cas9 ribonucleoprotein delivery (Paix *et al*. 2015). The resulting Bristol swap strains were named ECA401 *top-2*(*ean2[Q797M]*) and ECA402 *top-2*(*ean3[Q797M]*). The resulting Hawaii strains were named ECA547 *top-2*(*ean4[M797Q]*), ECA548 *top-2*(*ean5[M797Q]*), and ECA549 *top-2*(*ean6[M797Q]*).

---

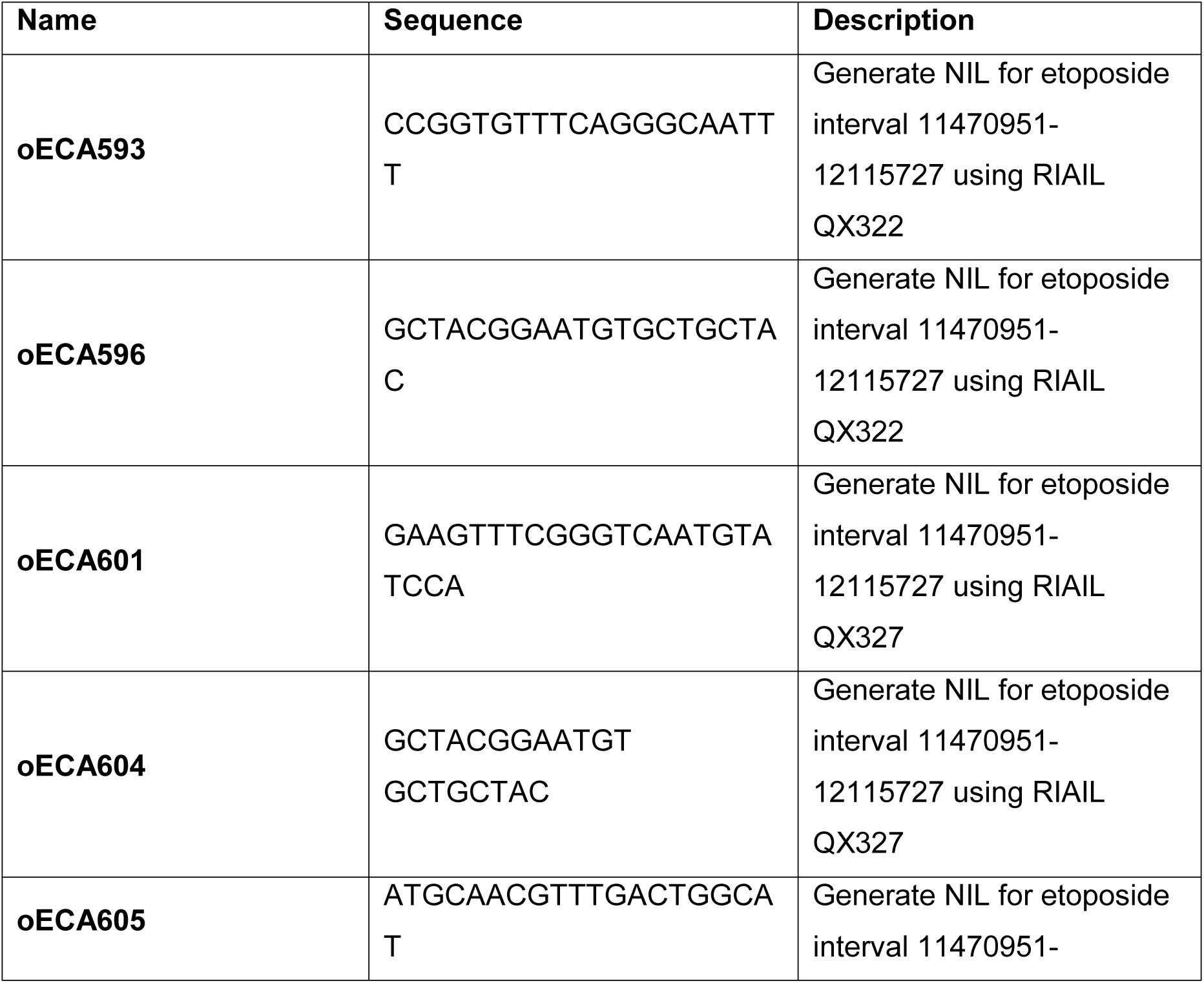
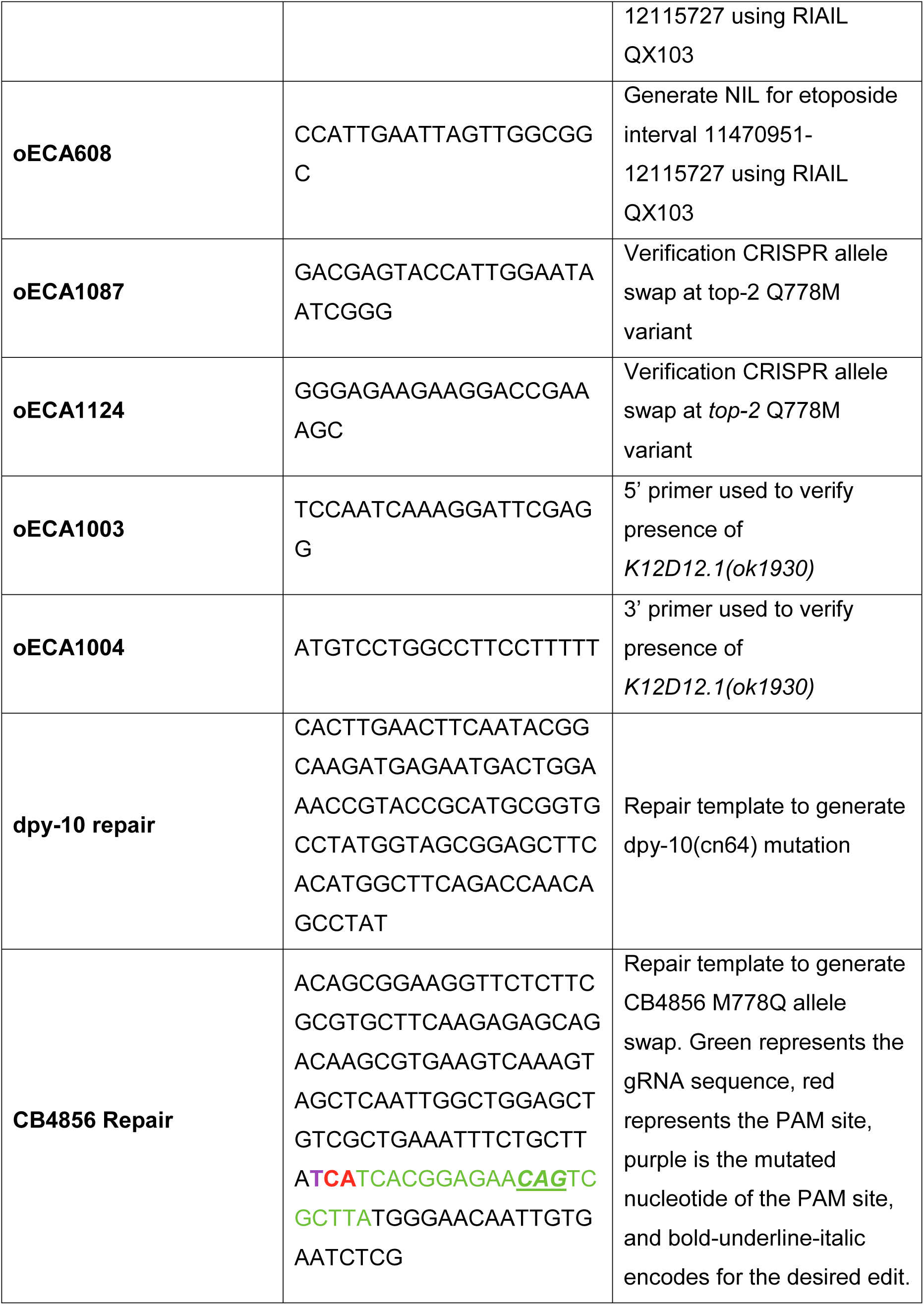
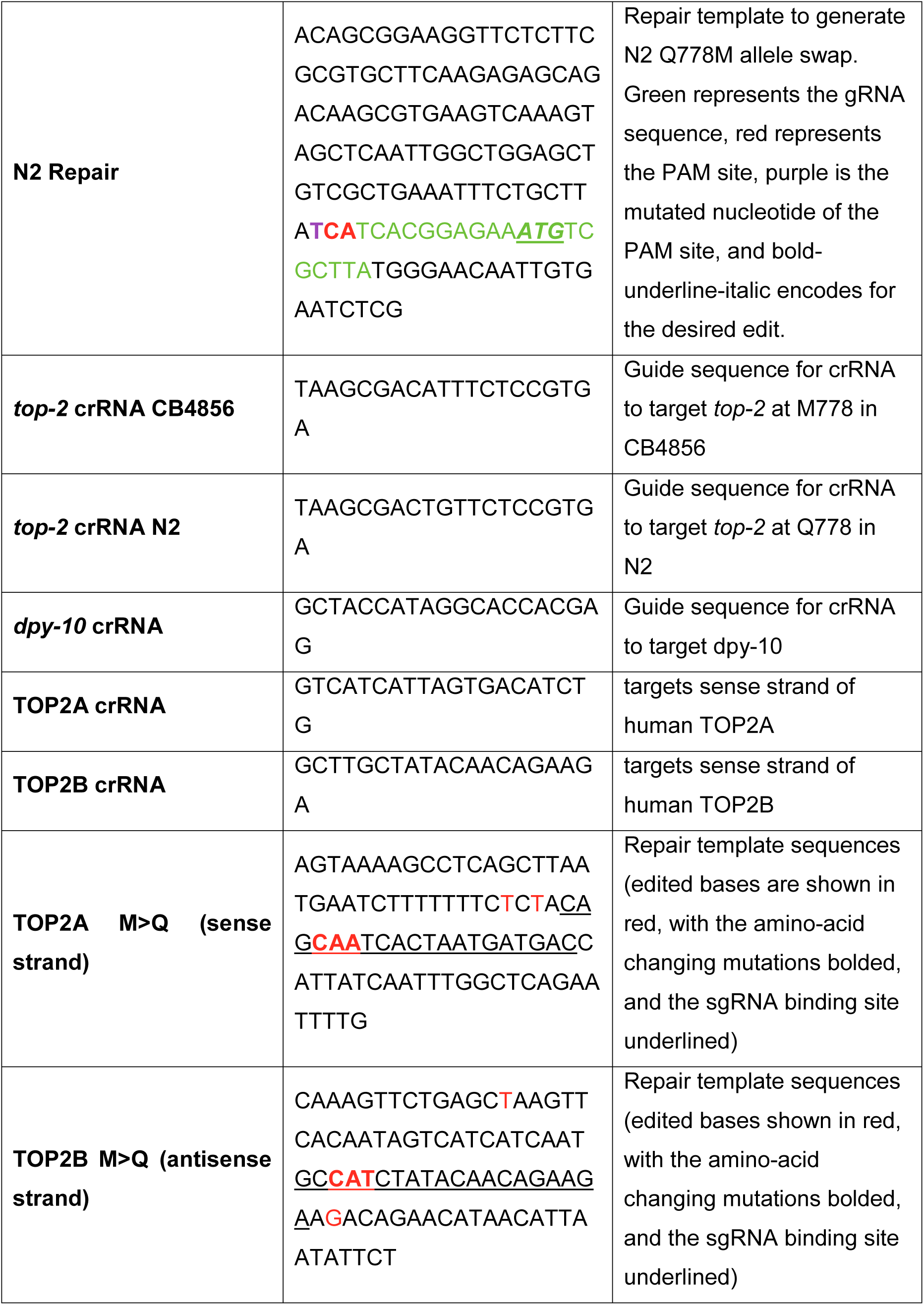
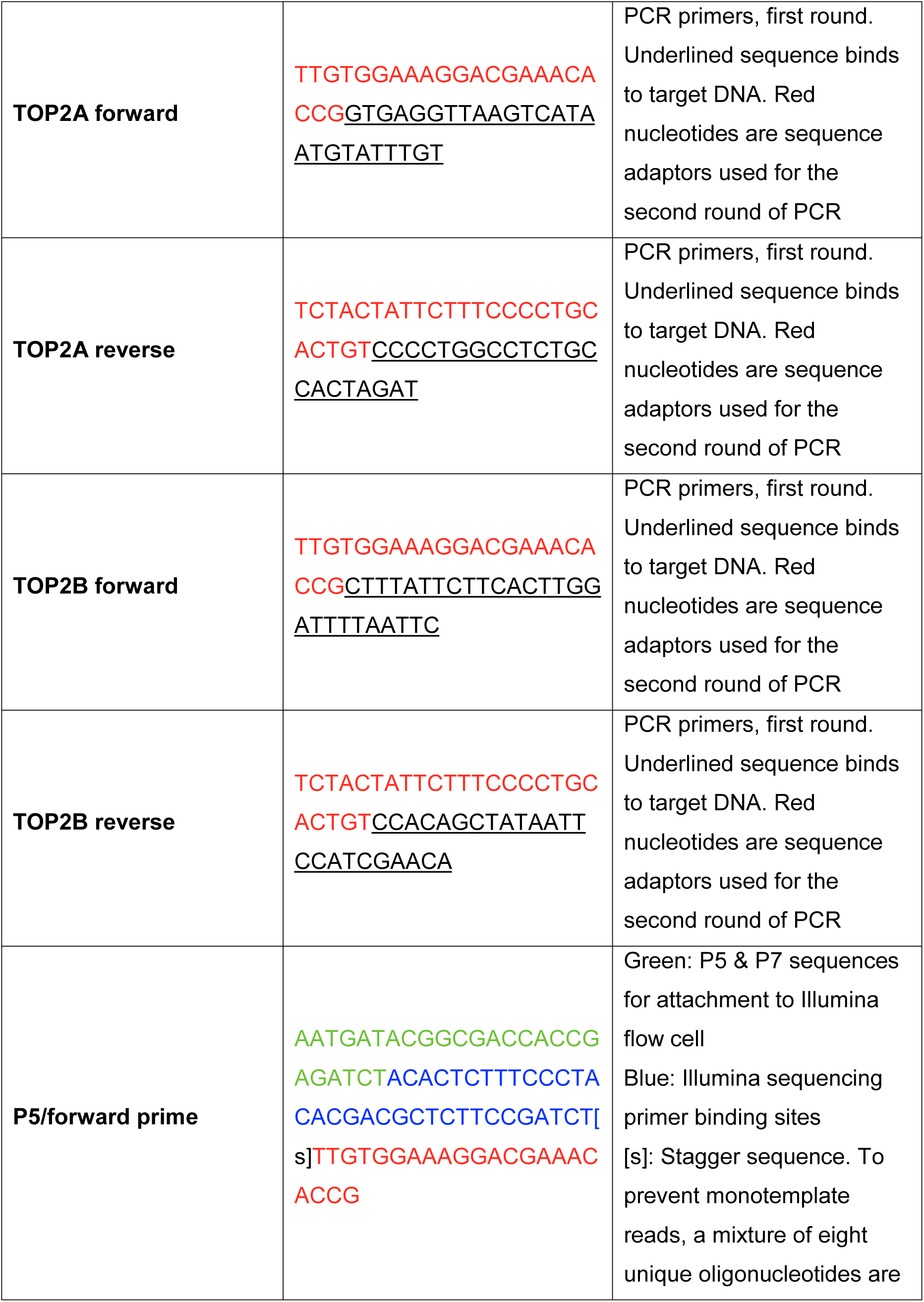
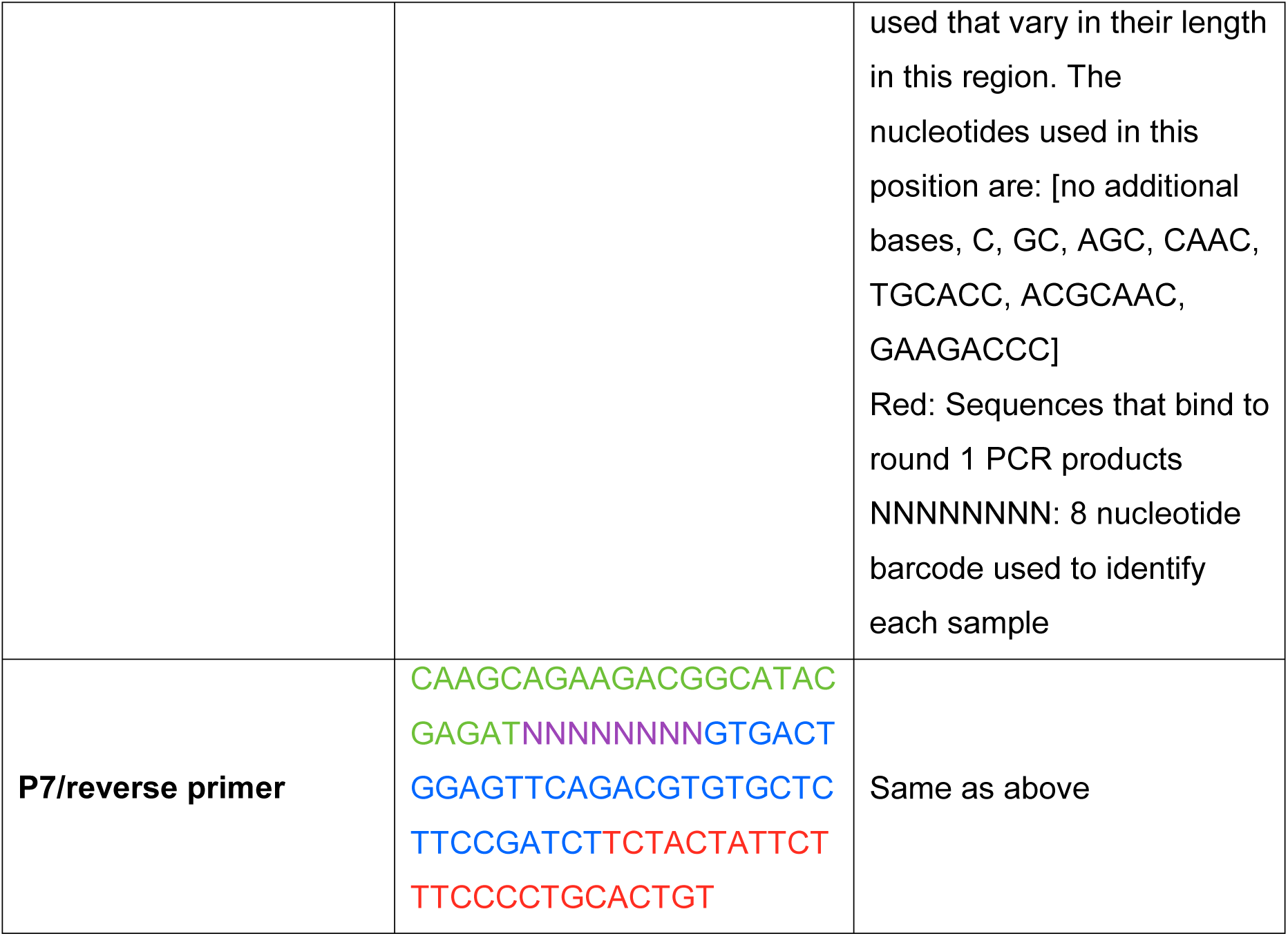
Primer Table

Strain table

**Strain table:**
| **Name** | **Allele Name** | **Genotype** |
| --- | --- | --- |
| **ECA215** | *eanIR135* | N2 (II:11.64 - 11.91 Mb) |
| **ECA216** | *eanIR136* | N2 (II:11.43 - 12.11 Mb) |
| **ECA219** | *eanIR139* | CB4856 (II:11.64 - 11.9 Mb) |
| **ECA220** | *eanIR140* | N2 (II:12.01 - 12.1 Mb) |
| **ECA338** | *K12D12.1(ok1930)* | (*K12D12.1(ok1930)/mIn1 [mIs14 dpy-10(e128)] II*) |
| **VC1474** | *K12D12.1(ok1930)* | (*K12D12.1(ok1930)/mIn1*<br>*[mIs14 dpy-10(e128)] II*) |
| **EG7952** | NA | [*oxTi207 [eft-3p::GFP::unc-54 3'UTR + hsp::peel-1 + NeoR + Cbr-unc-119(+)]*] |
| **ECA401** | *top-2(ean2[Q797M])* | N2 TOP2(Q797M) |
| **ECA402** | *top-2(ean3[Q797M])* | N2 TOP2(Q797M) |
| **ECA547** | *top-2(ean4[M797Q])* | CB4856 TOP2(M797Q) |
| **ECA548** | *top-2(ean5[M797Q])* | CB4856 TOP2(M797Q) |
| **ECA549** | *top-2(ean6[M797Q])* | CB4856 TOP2(M797Q) |

## References

Andersen, E.C. et al., 2015. A Powerful New Quantitative Genetics Platform, Combining Caenorhabditis elegans High-Throughput Fitness Assays with a Large Collection of Recombinant Strains. G3, 5(5), pp.g3.115.017178–920. Available at: http://g3journal.org/cgi/doi/10.1534/g3.115.017178.

Andersen, E.C. et al., 2014. A variant in the neuropeptide receptor npr-1 is a major determinant of Caenorhabditis elegans growth and physiology., 10(2), p.e1004156. Available at: http://dx.plos.org/10.1371/journal.pgen.1004156.

Andersen, E.C. et al., 2012. Chromosome-scale selective sweeps shape Caenorhabditis elegans genomic diversity. Nature genetics, 44(3), pp.285–290. Available at: http://dx.doi.org/10.1038/ng.1050.

Azarova, A.M. et al., 2007. Roles of DNA topoisomerase II isozymes in chemotherapy and secondary malignancies. Proceedings of the National Academy of Sciences of the United States of America, 104(26), pp.11014–11019. Available at: http://www.pnas.org/content/104/26/11014.full.

Bandele, O.J. & Osheroff, N., 2008. The efficacy of topoisomerase II-targeted anticancer agents reflects the persistence of drug-induced cleavage complexes in cells. Biochemistry, 47(45), pp.11900–11908. Available at: http://pubs.acs.org/doi/abs/10.1021/bi800981j.

Bloom, J.S. et al., 2013. Finding the sources of missing heritability in a yeast cross. Nature, 494(7436), pp.1–6. Available at: http://dx.doi.org/10.1038/nature11867.

Boddy, A.V., 2013. Genetics of cisplatin ototoxicity: confirming the unexplained? Clinical pharmacology and therapeutics, 94(2), pp.198–200. Available at: http://dx.doi.org/10.1038/clpt.2013.116.

Boyd, W.A., Smith, M.V. & Freedman, J.H., 2012. Caenorhabditis elegans as a model in developmental toxicology. Methods in molecular biology, 889(Chapter 3), pp.15–24. Available at: http://link.springer.com/10.1007/978-1-61779-867-2_3.

Brem, R.B. & Kruglyak, L., 2005. The landscape of genetic complexity across 5,700 gene expression traits in yeast. Proceedings of the National Academy of Sciences of the United States of America, 102(5), pp.1572–1577. Available at: http://www.pnas.org/content/102/5/1572.abstract.

Bromberg, K.D., Burgin, A.B. & Osheroff, N., 2003. A Two-drug Model for Etoposide Action against Human Topoisomerase IIα. The Journal of biological chemistry, 278(9), pp.7406–7412. Available at: http://www.jbc.org/content/278/9/7406.abstract.

Chen, S.H., Chan, N.-L. & Hsieh, T.-S., 2013. New mechanistic and functional insights into DNA topoisomerases. Annual review of biochemistry, 82(1), pp.139–170. Available at: http://www.annualreviews.org/doi/abs/10.1146/annurev-biochem-061809-100002.

Cingolani, P. et al., 2012. A program for annotating and predicting the effects of single nucleotide polymorphisms, SnpEff: SNPs in the genome of Drosophila melanogaster strain w1118; iso-2; iso-3. Fly, 6(2), pp.80–92. Available at: http://dx.doi.org/10.4161/fly.19695.

Cook, D.E., Zdraljevic, S., Roberts, J.P., et al., 2016. CeNDR, the Caenorhabditis elegans natural diversity resource. Nucleic acids research. Available at: http://dx.doi.org/10.1093/nar/gkw893.

Cook, D.E., Zdraljevic, S., Tanny, R.E., et al., 2016. The Genetic Basis of Natural Variation in Caenorhabditis elegans Telomere Length. Genetics, 204(1), pp.371–383. Available at: http://www.genetics.org/cgi/doi/10.1534/genetics.116.191148.

Cowell, I.G. et al., 2012. Model for MLL translocations in therapy-related leukemia involving topoisomerase IIβ-mediated DNA strand breaks and gene proximity. Proceedings of the National Academy of Sciences of the United States of America, 109(23), pp.8989–8994. Available at: http://dx.doi.org/10.1073/pnas.1204406109.

Davis, I.W. et al., 2007. MolProbity: all-atom contacts and structure validation for proteins and nucleic acids. Nucleic acids research, 35(Web Server issue), pp.W375–83. Available at: http://dx.doi.org/10.1093/nar/gkm216.

Demogines, A. et al., 2008. Identification and dissection of a complex DNA repair sensitivity phenotype in Baker’s yeast. PLoS genetics, 4(7), p.e1000123. Available at: http://dx.doi.org/10.1371/journal.pgen.1000123.

Deweese, J.E. & Osheroff, N., 2009. The DNA cleavage reaction of topoisomerase II: wolf in sheep’s clothing. Nucleic acids research, 37(3), pp.738–748. Available at: http://nar.oxfordjournals.org/lookup/doi/10.1093/nar/gkn937.

Ehrenreich, I.M. et al., 2010. Dissection of genetically complex traits with extremely large pools of yeast segregants. Nature, 464(7291), pp.1039–1042. Available at: http://www.nature.com/doifinder/10.1038/nature08923.

Endelman, J.B., 2011. Ridge Regression and Other Kernels for Genomic Selection with R Package rrBLUP. The Plant Genome Journal, 4(3), pp.250–256. Available at: https://www.crops.org/publications/tpg/abstracts/4/3/250.

Felix, C.A., Kolaris, C.P. & Osheroff, N., 2006. Topoisomerase II and the etiology of chromosomal translocations. DNA repair, 5(9-10), pp.1093–1108. Available at: http://dx.doi.org/10.1016/j.dnarep.2006.05.031.

Gao, H. et al., 1999. XK469, a selective topoisomerase IIβ poison. Proceedings of the National Academy of Sciences, 96(21), pp.12168–12173. Available at: http://www.pnas.org/content/96/21/12168.abstract.

Ghosh, R. et al., 2012. Natural Variation in a Chloride Channel Subunit Confers Avermectin Resistance in C. elegans. Science, 335(6068), pp.574–578. Available at: http://www.sciencemag.org/cgi/doi/10.1126/science.1214318.

Giacomini, K.M. et al., 2007. The pharmacogenetics research network: from SNP discovery to clinical drug response. Clinical pharmacology and therapeutics, 81(3), pp.328–345. Available at: http://dx.doi.org/10.1038/sj.clpt.6100087.

Gómez-Herreros, F. et al., 2013. TDP2-dependent non-homologous end-joining protects against topoisomerase II-induced DNA breaks and genome instability in cells and in vivo. PLoS genetics, 9(3), p.e1003226. Available at: http://dx.plos.org/10.1371/journal.pgen.1003226.

Huang, R.S. et al., 2007. A genome-wide approach to identify genetic variants that contribute to etoposide-induced cytotoxicity., 104(23), pp.9758–9763. Available at: http://www.pnas.org/content/104/23/9758.full.

Hunter, D.J., 2005. Gene-environment interactions in human diseases. Nature reviews. Genetics, 6(4), pp.287–298. Available at: http://dx.doi.org/10.1038/nrg1578.

Jacobson, M.P. et al., 2004. A hierarchical approach to all-atom protein loop prediction. Proteins, 55(2), pp.351–367. Available at: http://dx.doi.org/10.1002/prot.10613.

Jacobson, M.P. et al., 2002. On the role of the crystal environment in determining protein side-chain conformations. Journal of molecular biology, 320(3), pp.597–608. Available at: https://www.ncbi.nlm.nih.gov/pubmed/12096912.

Kang, H.M. et al., 2008. Efficient Control of Population Structure in Model Organism Association Mapping. Genetics, 178(3), pp.1709–1723. Available at: http://www.genetics.org/cgi/doi/10.1534/genetics.107.080101.

Kim, H. et al., 2014. A co-CRISPR strategy for efficient genome editing in Caenorhabditis elegans. Genetics, 197(4), pp.1069–1080. Available at: http://www.genetics.org/cgi/doi/10.1534/genetics.114.166389.

King, E.G. et al., 2014. Using Drosophila melanogaster to identify chemotherapy toxicity genes. Genetics, 198(1), pp.31–43. Available at: http://dx.doi.org/10.1534/genetics.114.161968.

Koba, M. & Konopa, J., 2005. [Actinomycin D and its mechanisms of action]. Postepy higieny i medycyny doswiadczalnej, 59, pp.290–298. Available at: https://www.ncbi.nlm.nih.gov/pubmed/15995596.

Koboldt, D.C. et al., 2013. The next-generation sequencing revolution and its impact on genomics. Cell, 155(1), pp.27–38. Available at: http://linkinghub.elsevier.com/retrieve/pii/S0092867413011410.

Lek, M. et al., 2016. Analysis of protein-coding genetic variation in 60,706 humans. Nature, 536(7616), pp.285–291. Available at: http://dx.doi.org/10.1038/nature19057.

Liti, G. & Louis, E.J., 2012. Advances in quantitative trait analysis in yeast. PLoS genetics, 8(8), p.e1002912. Available at: http://dx.doi.org/10.1371/journal.pgen.1002912.

Liu, J. et al., 2013. Identification of gene-environment interactions in cancer studies using penalization. Genomics, 102(4), pp.189–194. Available at: http://dx.doi.org/10.1016/j.ygeno.2013.08.006.

Low, S.-K. et al., 2013. Genome-wide association study of chemotherapeutic agent-induced severe neutropenia/leucopenia for patients in Biobank Japan. Cancer science, 104(8), pp.1074–1082. Available at: http://doi.wiley.com/10.1111/cas.12186.

Mariani, A. et al., 2015. Differential Targeting of Human Topoisomerase II Isoforms with Small Molecules. Journal of medicinal chemistry, 58(11), pp.4851–4856. Available at: http://dx.doi.org/10.1021/acs.jmedchem.5b00473.

Moen, E.L. et al., 2012. Pharmacogenomics of chemotherapeutic susceptibility and toxicity. Genome medicine, 4(11), p.90. Available at: http://dx.doi.org/10.1186/gm391.

Nitiss, J.L., 2009. Targeting DNA topoisomerase II in cancer chemotherapy. Nature reviews. Cancer, 9(5), pp.338–350. Available at: http://www.nature.com/doifinder/10.1038/nrc2607.

Paix, A. et al., 2015. High Efficiency, Homology-Directed Genome Editing in Caenorhabditis elegans Using CRISPR-Cas9 Ribonucleoprotein Complexes. Genetics, 201(1), pp.47–54. Available at: http://www.genetics.org/cgi/doi/10.1534/genetics.115.179382.

Park, J.-H. et al., 2012. Potential usefulness of single nucleotide polymorphisms to identify persons at high cancer risk: an evaluation of seven common cancers. Journal of clinical oncology: official journal of the American Society of Clinical Oncology, 30(17), pp.2157–2162. Available at: http://jco.ascopubs.org/cgi/doi/10.1200/JCO.2011.40.1943.

Perlstein, E.O. et al., 2007. Genetic basis of individual differences in the response to small-molecule drugs in yeast. Nature genetics, 39(4), pp.496–502. Available at: http://www.nature.com/doifinder/10.1038/ng1991.

Pommier, Y. et al., 2010. DNA Topoisomerases and Their Poisoning by Anticancer and Antibacterial Drugs. Chemistry & biology, 17(5), pp.421–433. Available at: http://dx.doi.org/10.1016/j.chembiol.2010.04.012.

Pommier, Y. et al., 1984. Formation and rejoining of deoxyribonucleic acid double-strand breaks induced in isolated cell nuclei by antineoplastic intercalating agents. Biochemistry, 23(14), pp.3194–3201. Available at: https://www.ncbi.nlm.nih.gov/pubmed/6087890.

Ratain, M.J. et al., 1987. Acute nonlymphocytic leukemia following etoposide and cisplatin combination chemotherapy for advanced non-small-cell carcinoma of the lung. Blood, 70(5), pp.1412–1417. Available at: https://www.ncbi.nlm.nih.gov/pubmed/2822173.

Rockman, M.V., Skrovanek, S.S. & Kruglyak, L., 2010. Selection at linked sites shapes heritable phenotypic variation in C. elegans. Science, 330(6002), pp.372–376. Available at: http://www.sciencemag.org/content/330/6002/372.long.

Sastry, G.M. et al., 2013. Protein and ligand preparation: parameters, protocols, and influence on virtual screening enrichments. Journal of computer-aided molecular design, 27(3), pp.221–234. Available at: http://dx.doi.org/10.1007/s10822-013-9644-8.

Schmidt, B.H., Osheroff, N. & Berger, J.M., 2012. Structure of a topoisomerase II-DNA-nucleotide complex reveals a new control mechanism for ATPase activity. Nature structural & molecular biology, 19(11), pp.1147–1154. Available at: http://www.nature.com/doifinder/10.1038/nsmb.2388.

Shimko, T.C. & Andersen, E.C., 2014. COPASutils: an R package for reading, processing, and visualizing data from COPAS large-particle flow cytometers. PloS one, 9(10), p.e111090. Available at: http://dx.plos.org/10.1371/journal.pone.0111090.

Sim, S.C., Altman, R.B. & Ingelman-Sundberg, M., 2011. Databases in the area of pharmacogenetics. Human mutation, 32(5), pp.526–531. Available at: http://dx.doi.org/10.1002/humu.21454.

Stern, D.L., 2014. Identification of loci that cause phenotypic variation in diverse species with the reciprocal hemizygosity test. Trends in genetics: TIG, 30(12), pp.547–554. Available at: http://dx.doi.org/10.1016/j.tig.2014.09.006.

Vejpongsa, P. & Yeh, E.T.H., 2014. Topoisomerase 2β: A Promising Molecular Target for Primary Prevention of Anthracycline-Induced Cardiotoxicity. Clinical Pharmacology & Therapeutics, 95(1), pp.45–52. Available at: http://onlinelibrary.wiley.com/doi/10.1038/clpt.2013.201/full.

Visscher, P.M. et al., 2012. Five years of GWAS discovery. American journal of human genetics, 90(1), pp.7–24. Available at: http://linkinghub.elsevier.com/retrieve/pii/S0002929711005337.

Wendorff, T.J. et al., 2012. The structure of DNA-bound human topoisomerase II alpha: conformational mechanisms for coordinating inter-subunit interactions with DNA cleavage. Journal of molecular biology, 424(3-4), pp.109–124. Available at: http://linkinghub.elsevier.com/retrieve/pii/S0022283612005815.

Willoughby, L.F. et al., 2013. An in vivo large-scale chemical screening platform using Drosophila for anti-cancer drug discovery. Disease models & mechanisms, 6(2), pp.521–529. Available at: http://dmm.biologists.org/cgi/doi/10.1242/dmm.009985.

Wu, C.-C. et al., 2013. On the structural basis and design guidelines for type II topoisomerase-targeting anticancer drugs. Nucleic acids research, 41(22), pp.10630–10640. Available at: http://nar.oxfordjournals.org/lookup/doi/10.1093/nar/gkt828.

Wu, C.-C. et al., 2011. Structural Basis of Type II Topoisomerase Inhibition by the Anticancer Drug Etoposide. Science, 333(6041), pp.456–459. Available at: http://www.sciencemag.org/cgi/doi/10.1126/science.1203963.

Yang, J. et al., 2009. Etoposide pathway. Pharmacogenetics and genomics, 19(7), pp.552–553. Available at: http://dx.doi.org/10.1097/FPC.0b013e32832e0e7f.

Yeh, E.T.H. & Bickford, C.L., 2009. Cardiovascular complications of cancer therapy: incidence, pathogenesis, diagnosis, and management. Journal of the American College of Cardiology, 53(24), pp.2231–2247. Available at: http://dx.doi.org/10.1016/j.jacc.2009.02.050.

Zhang, S. et al., 2012. Identification of the molecular basis of doxorubicin-induced cardiotoxicity. Nature medicine, 18(11), pp.1639–1642. Available at: http://dx.doi.org/10.1038/nm.2919.

